# Crosstalk between biochemical signaling and trafficking determines AMPAR dynamics in synaptic plasticity

**DOI:** 10.1101/2021.12.23.473965

**Authors:** M. K. Bell, P. Rangamani

## Abstract

Synaptic plasticity involves the modification of both biochemical and structural components of neurons. Many studies have revealed that the change in the number density of the glutamatergic receptor AMPAR at the synapse is proportional to synaptic weight update; increase in AMPAR corresponds to strengthening of synapses while decrease in AMPAR density weakens synaptic connections. The dynamics of AMPAR are thought to be regulated by upstream signaling, primarily the calcium-CaMKII pathway, trafficking to and from the synapse, and influx from extrasynaptic sources. Here, we have developed a set of models using compartmental ordinary differential equations to systematically investigate contributions of signaling and trafficking variations on AMPAR dynamics at the synaptic site. We find that the model properties including network architecture and parameters significantly affect the integration of fast upstream species by slower downstream species. Furthermore, we predict that the model outcome, as determined by bound AMPAR at the synaptic site, depends on (a) the choice of signaling model (bistable CaMKII or monostable CaMKII dynamics), (b) trafficking versus influx contributions, and (c) frequency of stimulus. Therefore, AMPAR dynamics can have unexpected dependencies when upstream signaling dynamics (such as CaMKII and PP1) are coupled with trafficking modalities.

## Introduction

A vast majority of the excitatory postsynaptic sites of synapses are housed in dendritic spines. Dendritic spines are small protrusions, 0.01 to 0.8 μm^3^ in volume, along the dendrites of neurons and act as signaling subcompartments, generating biochemical, electrical, and mechanical responses in response to stimuli from the presynaptic terminal [1]. Despite their small size, dendritic spines undergo complex biochemical signal transduction, spanning multiple timescales and pathways involving many protein kinases and phosphatases [2, 3]. Integration of these different pathways into a readout for learning and memory formation is the basis of synaptic plasticity, which refers to the ability of a synapse to regulate its connection strength through both biochemical and structural components [2].

A widely accepted and functional readout of synaptic plasticity is the density of the glutamatergic receptor, AMPAR (*α*-amino-3-hydroxy-5-methyl-4-isoxazolepropionic acid receptor), at the post-synaptic density (PSD) on the spine head [4]. Due to the importance of AMPAR for proper synaptic plasticity and neural function, dysregulation in either the underlying biochemical signaling or trafficking mechanisms of AMPAR can lead to severe consequences on learning, memory formation, and neural function, see Figure 1a.

**Figure 1:**
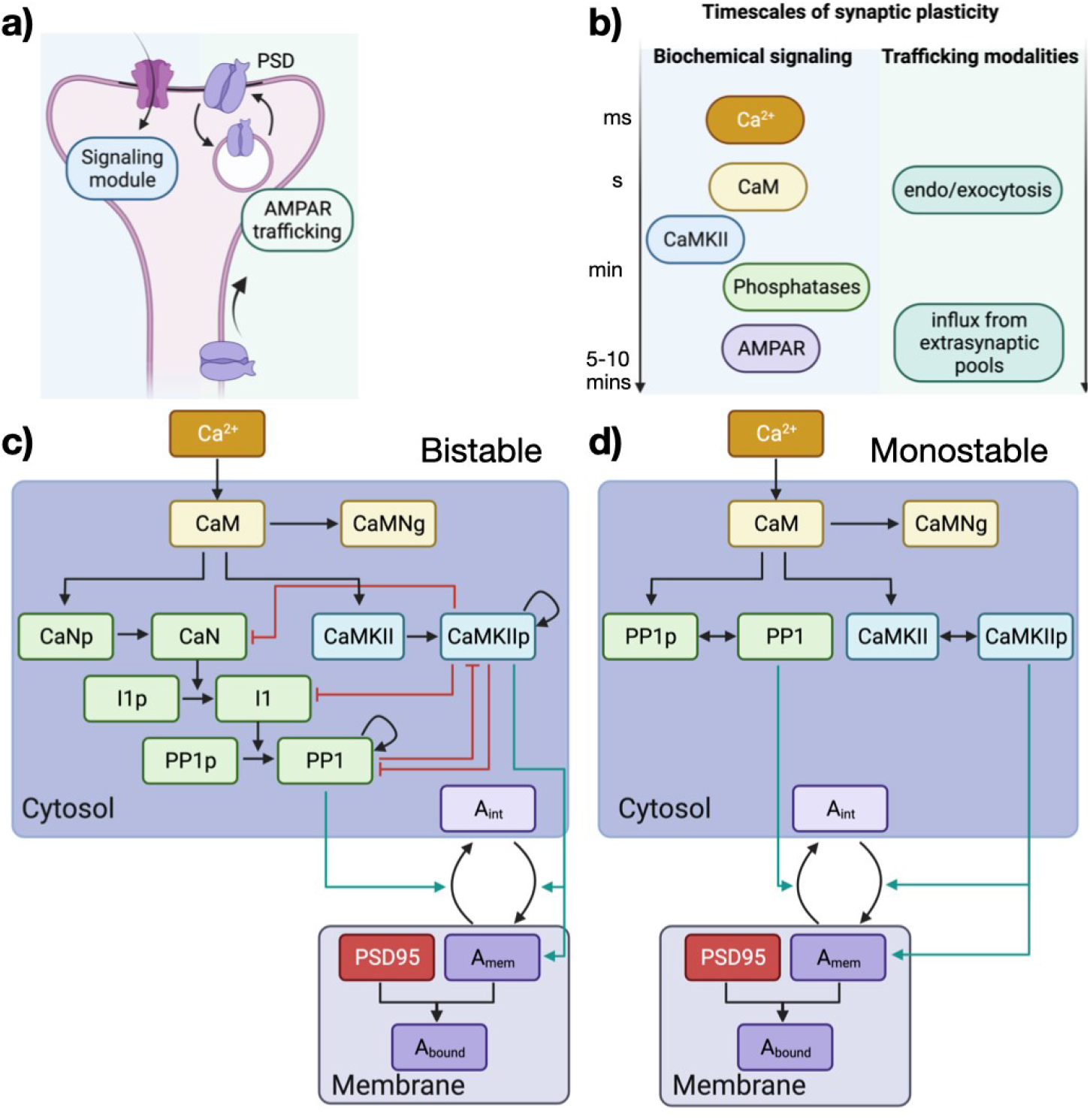
Schematic of AMPAR signaling and trafficking. a) AMPAR dynamics depend on a combination of signaling dynamics triggered through synaptic communication and movement of AMPAR through different trafficking mechanisms. b) Synaptic plasticity and specifically AMPAR modifications span several timescales from fast calcium influx to receptor trafficking into the dendritic spine. Although the signaling underlying synaptic plasticity is very complex, here we present two simplified biochemical signaling pathways to investigate the effects of both biochemical and trafficking variations on bound AMPAR at the PSD. The bistable model (c) and monostable model (d) represent two different mathematical formulations to describe how CaMKII and PP1 interact in dendritic spines. Black lines represent activation, red lines represent inhibition, and teal lines highlight biochemical influences on trafficking mechanisms. Made with Biorender.com.

Even though the processes associated with accumulation of AMPAR at the PSD are quite complex, to understand the events leading to the modification of AMPAR density on the spine head, we first provide a brief summary of the upstream signaling events organized by timescales, see Figure 1b for a schematic of these event timescales. The timescale of AMPAR increase/decrease at PSD after a stimulus varies from 10s [3, 5, 6] to a longer timescale of 60 minutes associated with long-term potentiation/depression (LTP/LTD) [7–9]. The main events occurring within these timescales are listed below.

- In response to a glutamate release event coupled with a voltage stimulus (primarily, an excitatory postsynaptic potential (EPSP) and a backpropagating action potential (BPAP)), N-methyl-D-aspartate receptors (NMDAR) and voltage sensitive calcium channels (VSCC) open on the spine membrane, resulting in an influx of Ca^2+^ into the spine. Calcium influx into the spine is the first step in numerous different signaling pathways important for synaptic function and from a modeling perspective is one of the most studied event in spines [10–16]. This voltage depolarization and subsequent calcium influx occurs over the millisecond timescale and is the fastest timescale considered in our model.
- Cytosolic Ca^2+^ rapidly binds a variety of different species over the millisecond to second timescale, notably calmodulin (CaM) [17], which in turn triggers various kinases and phosphatases including Calcium/calmodulin-dependent protein kinase II (CaMKII) and protein phosphatase 1 (PP1) [3, 18, 19]. Free cytosolic Ca^2+^ is also bound to various calcium buffers both in the cytoplasm and located on the membrane [10, 20]. CaM additionally binds to scaffolding proteins such as neurogranin (Ng) which impacts the available CaM for CaMKII activation [21].
- CaMKII is one of the most abundant proteins in the brain and its dynamics are vital for neural processes like synaptic plasticity, specifically LTP and LTD [22, 23]. CaMKII is known to increase the exocytosis rate of AMPAR to the surface of the spine head [6], promoting the idea that it is a molecular marker of memory [24, 25]. CaMKII is activated rapidly by Ca^2+^-bound CaM but its activity can remain elevated in the minute timescale due to its autophosphorylation dynamics (discussed more below) [26, 27]. CaMKII is also known to deactivate different phosphatases, including PP1 [28, 29]. In light of the critical role played by CaMKII in synaptic plasticity, a brief history of modeling is warranted. In [30], the idea that CaMKII activation exhibited bistability was proposed as a model to explain the long-time activation of CaMKII even after the stimulus was removed. In particular, [3] proposed a bistable model that included mutual inhibition and autoactivation of CaMKII and PP1 through Michaelis-Menten style dynamics [3, 30–32], see Figure 1c for a model of these bistable dynamics. This achieved multiple steady states of CaMKII and PP1 dependent on the level of Ca^2+^ input. While this system provided a switch that was protected from stochastic fluctuations and provided sustained CaMKII activity to model either LTP or LTD, experiments at the time only showed transient CaMKII activity [26, 33] and failed to find a hysteresis in the CaMKII-PP1 system [34, 35]. However, more recently, [36] experimentally showed the presence of CaMKII-PP1 hysteresis and bistability within individual holoenzymes in the presence of an NMDAR-derived peptide. They concluded that the CaMKII-PP1 system can act as a memory switch with bistability but only in the presence of NMDAR [36]. Other experiments showed that the observed bistability features required higher than relevant concentrations of Ca^2+^ suggesting that (a) bistability could exist at basal conditions if there are additional, currently unknown, signaling interactions that alter CaMKII dynamics [35], or (b) bistability might exist within localized domains such as within the PSD versus the spine cytoplasm [37], or (c) the observed hysteresis is not from true bistable dynamics but rather just a delayed return to a monostable state [36], which was supported by other models [38]. The temporal dynamics of CaMKII have also been extensively investigated experimentally. CaMKII was initially thought to have sustained activation because of its elevated levels following a brief stimulus and the necessity of CaMKII autophosphorylation for LTP [39]. However, more recent experiments show that CaMKII activation is actually transient and that LTP is associated with only transient CaMKII activity [33], see Figure 1d for a model of transient CaMKII and PP1 dynamics. Experimentally measured CaMKII dynamics are often fit with a biexponential decay with a fast and slow timescale [40]. While phosphastases are known to be involved in CaMKII inactivation, it was shown that phosphatases do not affect the fast decay dynamics of CaMKII [40]. Recently, we developed a multiscale model of CaMKII activation that considered monomer kinetics and holoenzyme kinetics and showed that CaMKII acts as a leaky integrator of Ca^2+^ pulses in the presence of Ng and PP1 [21], as seen experimentally in sLTP [41]. Another model also suggests that CaMKII is a leaky integrator of Ca^2+^ when in a PP1-rich environment [37].
- In addition to CaMKII, different phosphatases are also activated by Ca^2+^ -bound CaM and play important roles in signaling control loops for synaptic plasticity [28, 34, 42]. PP1 in particular is enriched in dendritic spines and is implicated in the regulation of LTD [19, 42, 43] and its activation is thought to increase the rate of endocytosis of AMPAR [30, 44]. PP1 has been hypothesized to deactivate CaMKII [28]; CaMKII is ultrasensitive in the presence of phosphatases and CaMKII within PSD was mainly dephosphorylated by PP1 [19]. Inhibiting protein phosphatases could convert short-term potentiation (STD) induced by exposure to K^+^ into LTP [33].
- CaMKII and PP1 influence AMPAR dynamics at the PSD by regulating exocytosis to the membrane and endocytosis from the membrane, respectively [44]. Specifically when CaMKII levels are high, AMPAR trafficking to the PSD is increased through exocytosis [45] and these conditions support the induction of LTP [25]. In contrast, when PP1 dominates the system, AMPAR density at the PSD decreases through endocytosis, supporting the induction of LTD [19, 46, 47]. Beyond endo- and exocytosis, there are other sources of AMPAR that can interact with AMPAR at the PSD. Pools of AMPAR exist on both the perisynaptic region and extrasynaptic regions both in the spine and on the dendrite, and laterally diffuse across the membrane into the PSD [48]. Synaptic activity, particularly CaMKII activation, can trigger movement of these AMPAR sources into the dendritic spine and towards the PSD region [23, 49, 50]. Therefore, AMPAR dynamics during this early synaptic plasticity phase occur over the 5-10 minute timescale [18]. Retention of the AMPAR at the PSD can occur through scaffolding molecules. TARPS (e.g. stargazin) binds to AMPAR very early on in the AMPAR lifecycle [49]. AMPAR also binds to scaffolding proteins, such as PSD95, to form bound AMPAR and this binding interaction plays a key role in determining AMPAR dynamics and localization in the PSD [22, 51].

Given the complexity of AMPAR density at the PSD, we seek to answer the following questions using computational modeling. How do the timescales of upstream biochemical signaling affect AMPAR dynamics? Specifically, how does the choice of biochemical signaling (bistable versus monostable model, Figure 1c versus d) of CaMKII/PP1 influence bound AMPAR dynamics? What is the role played by the trafficking modalities and influx of AMPAR on these dynamics? And finally, how do these response change with frequency of stimulus? To answer these questions, we developed a system of coupled compartmental ordinary differential equations. While we fully acknowledge that our model makes significant simplifications of the complex signaling processes[18], our focus is on understanding how these different timescales and contributions play an important role, see Figure 1. Using this model, we demonstrate that AMPAR dynamics depends in an expected manner on upstream signaling but unexpectedly on trafficking.

## Model development

To study the dynamics of AMPAR during synaptic plasticity, we constructed a system of compartmental deterministic ordinary differential equations (ODEs) that describes the influx of calcium, activation of CaMKII and PP1, and AMPAR dynamics on the membrane. We note that the pathways involved are quite complex and we make simplifying assumptions to probe the timescales of AMPAR dynamics on the membrane. Here we list these model assumptions, describe the key components of the model, and provide the governing equations.

### Model assumptions

- **Compartments:** We model a dendritic spine in the hippocampus as an two compartment system with a cytoplasm and plasma membrane. The cytoplasm has a volume of 0.06 μm^3^ based on the average dendritic spine volume [10, 52, 53]. The plasma membrane has a surface area of 0.8 μm^2^, which is the approximately surface area for an idealized spherical spine with a volume of 0.06 μm^3^ [10]. When converting between the volumetric cytoplasm and 2D plasma membrane compartments, we use a length scale conversion (n) of 0.1011 μm to convert units [54]. All species are well mixed within their compartment. We model the spine as an isolated compartment besides the initial Ca^2+^ influx and the influx of CaMKII-dependent membrane AMPAR [26, 55].
- **Timescales:** We focus on AMPAR dynamics associated with synaptic plasticity on the 10 minute timescale [18].
- **Reaction types:** A vast majority of the reactions are modeled as either mass action dynamics or enzymatic Hill function reactions (See Tables 1, 2, 3, 4, 5, 6). Several components, including the dynamics of NMDAR and VSCC, are custom equations that describe how their activation depends on voltage [10, 56]. AMPAR endocytosis and exocytosis are taken as modified mass action dynamics with additional dependence on activated CaMKII and PP1 [3, 30]. AMPAR influx from extrasynaptic pools outside of the dendritic spine is modeled as a time-dependent and activated CaMKII-dependent process to represent activity-dependent diffusion of membrane AMPAR into the system [50, 57]. This AMPAR influx term means that the system does not have mass conservation for the membrane AMPAR (Amem) species.
- **Kinetic parameters:** Reaction rates were taken from previous studies [10, 28] or approximately fit to experimental data. In particular, the rates for CaMKII and PP1 decay dynamics for the monostable model were fit to experimental data [12, 40, 41], see Figure 2j-k.

**Figure 2:**
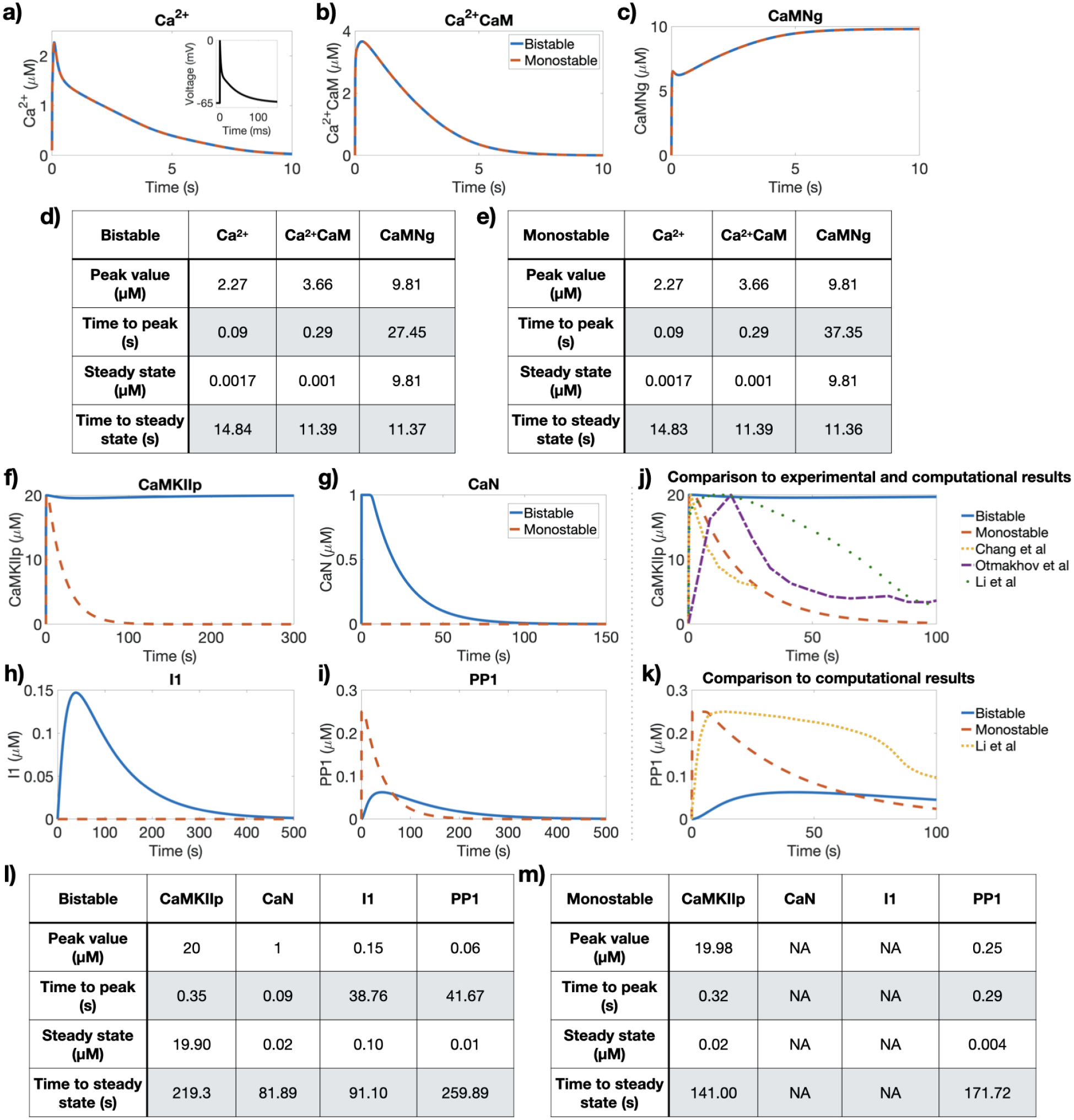
Early and intermediate timescale events. Temporal dynamics of cytosolic Ca^2+^ (a), activated Ca^2+^/CaM complex (b), and CaM/Ng complex (c) all show similar dynamics for both the bistable (blue line) and monostable models (red line). Black inset in (a) is the voltage stimulus that represents a EPSP and BPAP separated by 2 ms. Peak and steady state values and times for Ca^2+^, Ca^2+^/CaM complex, and CaM/Ng complex for the bistable (d) and monostable (e) models show similar dynamics for early timescale events. Temporal dynamics of activated CaMKII (f), activated CaN (g), activated I1 (h), and activated PP1 (i) for both the bistable and monostable models show very different results. Note that the monostable model does not involve CaN and I1. j) Comparison of CaMKII dynamics between the bistable (blue) and monostable (red) models, experimental observations [40, 41], and other models [12]. k) Comparison of PP1 dynamics between both the bistable (blue) and monostable (red) models and another model [12]. Peak and steady state values and times for activated CaMKII, activated CaN, activated I1, and activated PP1 for the bistable (l) and monostable (m) models show different dynamics.

**Table 1:** AMPAR pathway reactions, reaction types, and reaction rates used in the bistable model

|  | List of Reactions | Reaction Type | Reaction Rate |
| --- | --- | --- | --- |
| Module 0: | Ca<sup>2+</sup> Influx |  |  |
| 0 | $\text{Ca}_{\text{ECS}}^{2+} \rightleftharpoons \text{Ca}_{\text{cyto}}^{2+}$ | Custom<sup>(0)</sup> | $R_0 = J_{\text{influx}} - J_{\text{efflux}}$ |
| Module 1: | Ca<sup>2+</sup>/CaM Complex Formation |  |  |
| 1 | $3 \text{Ca}^{2+} + \text{CaM} \xrightleftharpoons[k_{b1}]{k_{f1}} \text{Ca}^{2+}\text{CaM}$ | MA<sup>(1)</sup> | $R_1 = k_{f1}[\text{Ca}^{2+}]^3[\text{CaM}] - k_{b1}[\text{Ca}^{2+}\text{CaM}]$ |
| Module 2: | CaM/Ng Complex Formation |  |  |
| 2 | $\text{CaM} + \text{Ng} \xrightleftharpoons[k_{b2}]{k_{f2}} \text{CaMNg}$ | MA | $R_2 = k_{f2}[\text{CaM}][\text{Ng}] - k_{b2}[\text{CaMNg}]$ |
| Module 3: | CaMKII activation and deactivation |  |  |
| 3 | $\text{CaMKII} + \text{Ca}^{2+}\text{CaM} \xrightarrow{\text{Kcat}_3} \text{CaMKIIP}$ | CB<sup>(2,3)</sup> | $R_3 = \frac{\text{Kcat}_3[\text{Ca}^{2+}\text{CaM}]^4[\text{CaMKII}]}{K_{m3}^4 + [\text{Ca}^{2+}\text{CaM}]^4}$ |
| 4 | $\text{CaMKII} + \text{CaMKIIP} \xrightarrow{\text{Kcat}_4} \text{CaMKIIP}$ | CB | $R_4 = \frac{\text{Kcat}_4[\text{CaMKIIP}][\text{CaMKII}]}{K_{m4} + [\text{CaMKIIP}]}$ |
| 5 | $\text{CaMKIIP} + \text{PP1} \xrightarrow{\text{Kcat}_5} \text{CaMKII}$ | CB | $R_5 = \frac{\text{Kcat}_5[\text{PP1}][\text{CaMKIIP}]}{K_{m5} + [\text{CaMKIIP}]}$ |
| Module 4: | Phosphatase cascade activation and deactivation |  |  |
| 6 | $\text{CaNp} + \text{Ca}^{2+}\text{CaM} \xrightarrow{\text{Kcat}_6} \text{CaN}$ | CB | $R_6 = \frac{\text{Kcat}_6[\text{Ca}^{2+}\text{CaM}]^4[\text{CaNp}]}{K_{m6}^4 + [\text{Ca}^{2+}\text{CaM}]^4}$ |
| 7 | $\text{CaN} + \text{CaMKIIP} \xrightarrow{\text{Kcat}_7} \text{CaNp}$ | CB | $R_7 = \frac{\text{Kcat}_7[\text{CaMKIIP}][\text{CaN}]}{K_{m7} + [\text{CaMKIIP}]}$ |
| 8 | $\text{I}_1\text{p} + \text{CaN} \xrightarrow{\text{Kcat}_8} \text{I}_1$ | CB | $R_8 = \frac{\text{Kcat}_8[\text{CaN}][\text{I}_1\text{p}]}{K_{m8} + [\text{I}_1\text{p}]}$ |
| 9 | $\text{I}_1 + \text{CaMKIIP} \xrightarrow{\text{Kcat}_9} \text{I}_1\text{p}$ | CB | $R_9 = \frac{\text{Kcat}_9[\text{CaMKIIP}][\text{I}_1]}{K_{m9} + [\text{I}_1]}$ |
| 10 | $\text{PP1p} + \text{I1} \xrightarrow{\text{Kcat}_{10}} \text{PP1}$ | CB | $R_{10} = \frac{\text{Kcat}_{10}[\text{I1}][\text{PP1p}]}{K_{m10} + [\text{PP1p}]}$ |
| 11 | $\text{PP1p} + \text{PP1} \xrightarrow{\text{Kcat}_{11}} \text{PP1}$ | CB | $R_{11} = \frac{\text{Kcat}_{11}[\text{PP1}][\text{PP1p}]}{K_{m11} + [\text{PP1p}]}$ |
| 12 | $\text{PP1} + \text{CaMKIIP} \xrightarrow{\text{Kcat}_{12}} \text{PP1p}$ | CB | $R_{12} = \frac{\text{Kcat}_{12}[\text{CaMKIIP}][\text{PP1}]}{K_{m12} + [\text{PP1}]}$ |
| Module 5: | AMPA trafficking |  |  |
| 13 | $\text{Aint} \xrightleftharpoons[k_{b13}(\text{PP1})]{k_{f13}(\text{CaMKIIP})} \text{Amem}$ | MA | $R_{13} = (c_1[\text{CaMKIIP}] + c_3)[\text{Aint}] - (c_2[\text{PP1}] + c_4)[\text{Amem}]$ |
| 14 | $\emptyset \xrightarrow{k_{f14}(\text{CaMKIIP})} \text{Amem}$ | Custom<sup>(4)</sup> | $R_{14} = (k_{f14}(t))[\text{CaMKIIP}]$ |
| Module 6: | AMPA/Scaffolding Complex |  |  |
| 15 | $\text{Amem} + \text{PSD}_{95} \xrightleftharpoons[k_{b15}]{k_{f15}} \text{ABound}$ | MA | $R_{15} = k_{f15}[\text{Amem}][\text{PSD}_{95}] - k_{b15}[\text{ABound}]$ |

1 <sup>(0)</sup> Custom taken from [10]: see Supplemental material for details. $J_{\text{influx}} = J_{\text{NMDAR}} + J_{\text{VSCC}} + J_{\text{leak}}$ and $J_{\text{efflux}} = J_{\text{PMCA}} + J_{\text{NCX}}$ ; <sup>(1)</sup> MA: Mass Action; <sup>(2)</sup> CB: Cooperative Binding; <sup>(3)</sup> Enzyme is in bold type. <sup>(4)</sup> Custom is given in Supplemental material.

**Table 2:**
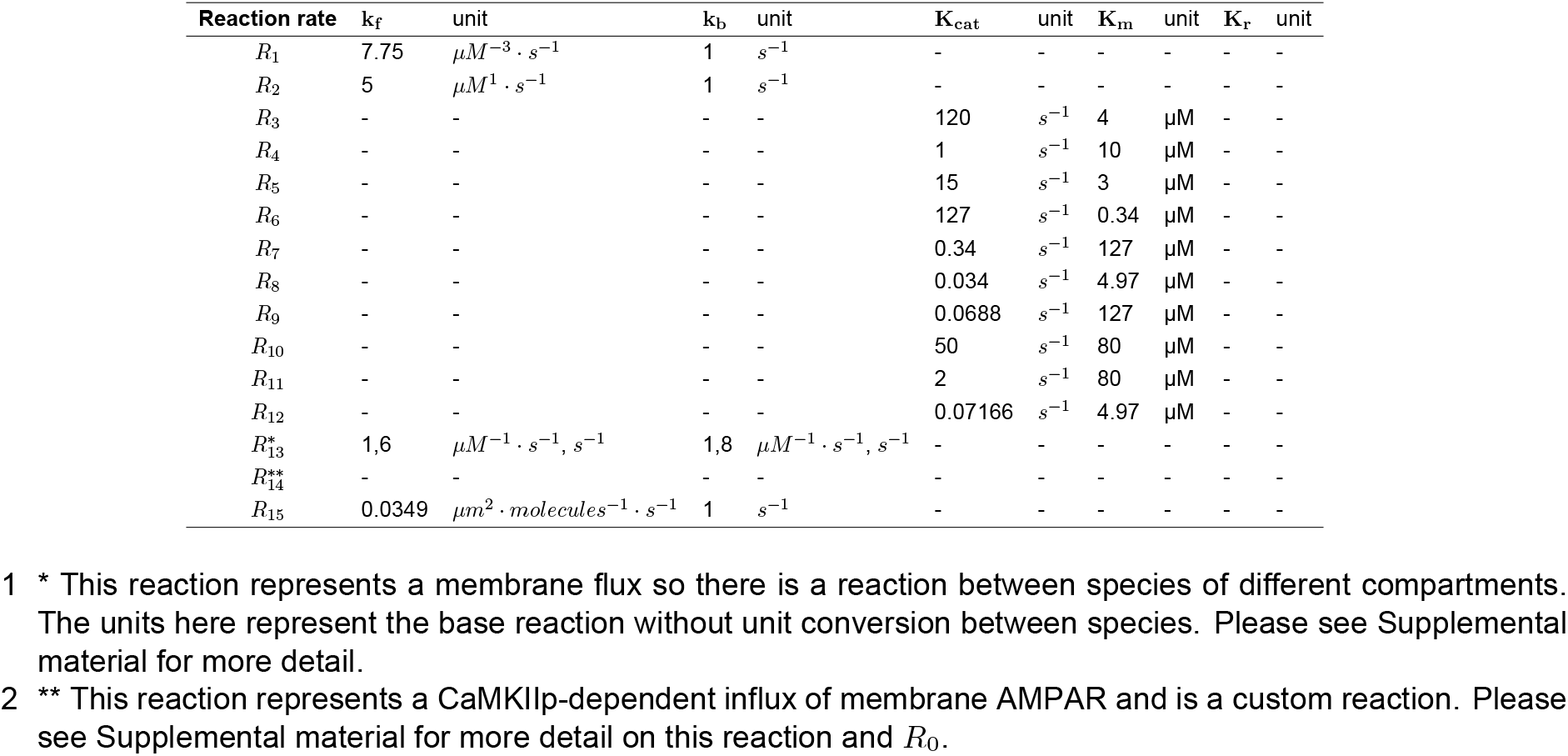
Reaction parameters calculated for the bistable model

**Table 3:** Initial concentrations and ordinary differential equations for bistable model

| Species | Initial condition | Units | ODE |
| --- | --- | --- | --- |
| $Ca^{2+}$ | 0.1 | $\mu M$ | $\frac{d[Ca^{2+}]}{dt} = R_0 - 3R_1$ |
| CaM | 10 | $\mu M$ | $\frac{d[CaM]}{dt} = -R_1$ |
| $Ca^{2+}CaM$ | 0 | $\mu M$ | $\frac{d[Ca^{2+}CaM]}{dt} = R_1$ |
| Ng | 20 | $\mu M$ | $\frac{d[Ng]}{dt} = -R_2$ |
| CaMNg | 0 | $\mu M$ | $\frac{d[CaMNg]}{dt} = R_2$ |
| CaMKII | 20<sup>1</sup> | $\mu M$ | $\frac{d[CaMKII]}{dt} = -R_3 - R_4 + R_5$ |
| CaMKIIp | 0 | $\mu M$ | $\frac{d[CaMKIIp]}{dt} = R_3 + R_4 - R_5$ |
| CaNp | 1 | $\mu M$ | $\frac{d[CaNp]}{dt} = -R_6 + R_7$ |
| CaN | 0 | $\mu M$ | $\frac{d[CaN]}{dt} = R_6 - R_7$ |
| I1p | 1.8 | $\mu M$ | $\frac{d[I1p]}{dt} = -R_8 + R_9$ |
| I1 | 0 | $\mu M$ | $\frac{d[I1]}{dt} = R_8 - R_9$ |
| PP1p | 0.25<sup>1</sup> | $\mu M$ | $\frac{d[PP1p]}{dt} = -R_{10} - R_{11} + R_{12}$ |
| PP1 | 0 | $\mu M$ | $\frac{d[PP1]}{dt} = R_{10} + R_{11} - R_{12}$ |
| Aint | 0.1597 | $\mu M$ | $\frac{d[Aint]}{dt} = -R_{13}$ |
| Amem | 7.288 | $\frac{molecules}{\mu m^2}$ | $\frac{d[Amem]}{dt} = R_{13} + R_{14} - R_{15}$ |
| PSD95 | 4000 | $\frac{molecules}{\mu m^2}$ | $\frac{d[PSD95]}{dt} = -R_{15}$ |
| ABound | 1000 | $\frac{molecules}{\mu m^2}$ | $\frac{d[ABound]}{dt} = R_{15}$ |

<sup>1</sup> Initial condition is varied in some of the simulations.
In simulations, membrane components are converted to moles per meter squared.

**Table 4:** AMPAR pathway reactions, reaction types, and reaction rates used in the monostable model

|  | List of Reactions | Reaction Type | Reaction Rate |
| --- | --- | --- | --- |
| Module 0: | Ca<sup>2+</sup> Influx |  |  |
| 0 | Ca<sub>ECS</sub><sup>2+</sup> ⇌ Ca<sub>cyto</sub><sup>2+</sup> | Custom<sup>(0)</sup> | $R_0 = J_{influx} - J_{efflux}$ |
| Module 1: | Ca<sup>2+</sup>/CaM Complex Formation |  |  |
| 1 | 3 Ca<sup>2+</sup> + CaM $\xrightleftharpoons[k_{b1}]{k_{f1}}$ Ca<sup>2+</sup>CaM | MA<sup>(1)</sup> | $R_1 = k_{f1}[Ca^{2+}]^3[CaM] - k_{b1}[Ca^{2+}CaM]$ |
| Module 2: | CaM/Ng Complex Formation |  |  |
| 2 | CaM + Ng $\xrightleftharpoons[k_{b2}]{k_{f2}}$ CaMNg | MA | $R_2 = k_{f2}[CaM][Ng] - k_{b2}[CaMNg]$ |
| Module 3: | CaMKII activation and deactivation |  |  |
| 3 | CaMKII + **Ca<sup>2+</sup>CaM** $\xrightarrow{K_{cat3}}$ CaMKIIP | CB<sup>(2,3)</sup> | $R_3 = \frac{K_{cat3}[Ca^{2+}CaM]^4[CaMKII]}{K_{m3}^4 + [Ca^{2+}CaM]^4}$ |
| 4 | CaMKIIP $\xrightarrow{k_{f4}}$ CaMKII | MA | $R_4 = k_{f4}[CaMKIIP]$ |
| Module 4: | Phosphatase cascade activation and deactivation |  |  |
| 5 | PP<sub>1</sub>P + **Ca<sup>2+</sup>CaM** $\xrightarrow{K_{cat5}}$ PP<sub>1</sub> | CB | $R_5 = \frac{K_{cat5}[Ca^{2+}CaM]^4[PP_1P]}{K_{m5}^4 + [Ca^{2+}CaM]^4}$ |
| 6 | PP<sub>1</sub> $\xrightarrow{k_{f6}}$ PP<sub>1</sub>P | MA | $R_6 = k_{f6}[PP_1]$ |
| Module 5: | AMPA trafficking |  |  |
| 7 | Aint $\xrightleftharpoons[k_{b7}(PP_1)]{k_{f7}(CaMKIIP)}$ Amem | MA | $R_7 = (c_1[CaMKIIP] + c_3)[Aint] - (c_2[PP_1] + c_4)[Amem]$ |
| 8 | $\emptyset \xrightarrow{k_{f8}(CaMKIIP)}$ Amem | Custom<sup>(4)</sup> | $R_8 = (k_{f8}(t)[CaMKIIP])$ |
| Module 6: | AMPA/Scaffolding Complex |  |  |
| 9 | Amem + PSD<sub>95</sub> $\xrightleftharpoons[k_{b9}]{k_{f9}}$ ABound | MA | $R_9 = k_{f9}[Amem][PSD95] - k_{b9}[ABound]$ |

- 1 <sup>(0)</sup> Custom taken from [10]: see Supplemental material for details. $J_{influx} = J_{NMDAR} + J_{VSCC} + J_{leak}$ and $J_{efflux} = J_{PMCA} + J_{NCX}$ ; <sup>(1)</sup> MA: Mass Action; <sup>(2)</sup> CB: Cooperative Binding; <sup>(3)</sup> Enzyme is in bold type. <sup>(4)</sup> Custom is given in Supplemental material.

**Table 5:**
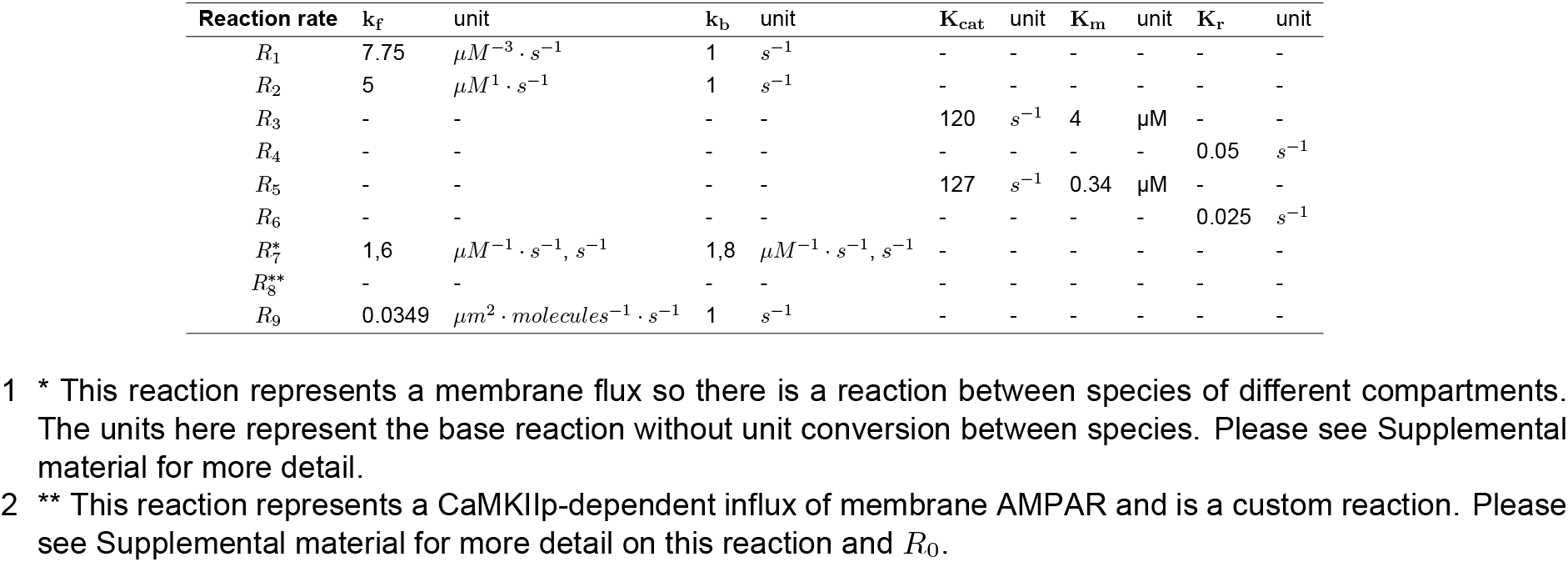
Reaction parameters calculated for the monostable model

**Table 6:** Initial concentrations and ordinary differential equations for bistable model

| Species | Initial condition | Units | ODE |
| --- | --- | --- | --- |
| Ca<sup>2+</sup> | 0.1 | μM | $\frac{d[Ca^{2+}]}{dt} = R_0 - 3R_1$ |
| CaM | 10 | μM | $\frac{d[CaM]}{dt} = -R_1$ |
| Ca<sup>2+</sup>CaM | 0 | μM | $\frac{d[Ca^{2+}CaM]}{dt} = R_1$ |
| Ng | 20 | μM | $\frac{d[Ng]}{dt} = -R_2$ |
| CaMNg | 0 | μM | $\frac{d[CaMNg]}{dt} = R_2$ |
| CaMKII | 20<sup>1</sup> | μM | $\frac{d[CaMKII]}{dt} = -R_3 + R_4$ |
| CaMKIIP | 0 | μM | $\frac{d[CaMKIIP]}{dt} = R_3 - R_4$ |
| PP1p | 0.25<sup>1</sup> | μM | $\frac{d[PP1p]}{dt} = -R_5 + R_6$ |
| PP1 | 0 | μM | $\frac{d[PP1]}{dt} = R_5 - R_6$ |
| Aint | 0.1597 | μM | $\frac{d[Aint]}{dt} = -R_7$ |
| Amem | 7.288 | $\frac{\text{molecules}}{\mu\text{m}^2}$ | $\frac{d[Amem]}{dt} = R_7 + R_8 - R_9$ |
| PSD95 | 4000 | $\frac{\text{molecules}}{\mu\text{m}^2}$ | $\frac{d[PSD95]}{dt} = -R_9$ |
| ABound | 1000 | $\frac{\text{molecules}}{\mu\text{m}^2}$ | $\frac{d[ABound]}{dt} = R_9$ |

<sup>1</sup> Initial condition is varied in some of the simulations.
In simulations, membrane components are converted to moles per meter squared.

### Key components

The goal of this model is to investigate AMPAR dynamics during spine activation and the key upstream kinase and phosphatase activity. We highlight the key model components below.

- **Stimulus and Ca^2+^ influx:** We simulate the activation of a dendritic spine due to voltage depolarization so the model stimulus is a time dependence voltage profile representing an EPSP and a BPAP offset by 2 ms [10]. The voltage depolarization is assumed to accompany a glutamate release, such that both NMDARs and VSCCs are activated, allowing for calcium influx into the spine. We do not explicitly model the glutamate release from the presynapse or the diffusion of glutamate in the extracellular space. We also do not explicitly model extracellular calcium, as we assume it has such a high concentration that the calcium influx into the dendritic spine has negligible effects [10]. After Ca^2+^ floods into the spine through NMDAR and VSCC, it is pumped out of the spine through PM Ca^2+^-ATPase (PMCA) pumps and sodium–calcium exchanger (NCX) pumps [10]. Ca^2+^ influx into the spine is also rapidly buffered by downstream signaling species, and here we explicitly model Ca^2+^ binding to Calmodulin (CaM) [28]. We do not consider additional buffers here due to explicit modeling of binding to CaM.
- **Calmodulin activation:** Calmodulin (CaM) is quickly bound to Ca^2+^ in dendritic spines and we simplify Ca^2+^/CaM binding in a single mass action equation. We note that Ca^2+^/CaM binding is a multistate process with different intermediates [21, 58]; we simplify the process to focus on key timescales of CaM. We denote the activated CaM complex (CaM bound to 4 Ca^2+^, Ca^2+^_4_CaM) as Ca^2+^CaM.
- **Activation of downstream components by calmodulin:** CaM acts as a key propagating signal by binding with the scaffolding protein neurogranin (Ng) [21], and activated CaM activates both CaMKII and the phosphatase cascade [28].
- **CaMKII and phosphatase dynamics:** We investigate two different models for CaMKII and phosphatase dynamics, shown in Fig. 1c and d.
  – **Bistable model:** In the bistable model, activated CaM triggers the phosphorylation of CaMKII, which can then autophosphorylate itself [3, 28, 31, 59]. Activated CaM triggers the dephosphorylation of calneurin (CaN), which is the first step in a phosphatase cascade involving inhibitor-1 (I1) and PP1, where CaN activates I1 and I1 activates PP1 [28, 60], see Supplemental Material for a comment on I1. Activated PP1 can then autodephosphorylate itself [28, 61]. Active CaMKII can deactivate each of the active phosphatases. Activated PP1 can dephosphorylate CaMKII, while activated CaMKII can phosphorylate PP1, so the active species of both CaMKII and PP1 deactivate the other species [36, 61]. We refer to this model as the bistable model because this system can achieve different steady state behavior of CaMKII and PP1 dependent on different stimulus, initial conditions, and other factors [3].
  – **Monostable model:** In the monostable model, activated CaM triggers the phosphorylation of CaMKII, but CaMKII deactivates over time proportionally to its activated concentration [26, 62]. Similarly, activated CaM triggers the activation of PP1, which then decays to its inactive state proportionally to its active concentration. We refer to this model as the monostable model because the rate of decrease for both active CaMKII and active PP1 is linearly proportional to their concentrations and therefore only produce a single steady state.
- **AMPAR trafficking and signaling:** AMPAR is a membrane-bound protein; it is located on the plasma membrane and on endosomal membranes, and undergoes both basal and activity dependent endocytosis and exocytosis [63, 64]. In our model, we incorporate CaMKII-dependent exocytosis [45] and PP1-dependent endocytosis [3]. AMPARs also diffuses into the spine in response to CaMKII activity through lateral membrane diffusion from extrasynaptic pools on the dendrite [45, 48]. On the membrane, AMPAR binds to scaffolding proteins, such as PSD95, to stabilize at the PSD [65]. We assume that there is a pool of PSD95 available to bind with membrane AMPAR. This bound AMPAR/PSD95 complex, denoted as bound AMPAR, is the model readout.

### Governing equations

We construct a compartmental system of ordinary differential equations that represent signaling species in the two different compartments, the cytoplasm and the plasma membrane. The temporal dynamics of each signaling species, c_i_, is given by

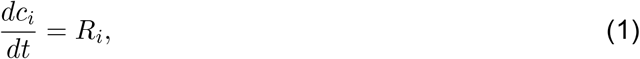

where *R_i_* represents the net flux of the species *i*. The reactions governing the dynamics of each species are the sum of sources and sinks such that *R_i_* = *R_i,source_* – *R_i,sink_*. For membrane AMPAR, this coupled system can be understood through a series of CaMKII and PP1 dependent sources and sinks for AMPAR on the membrane. Unbound AMPAR on the membrane increases through CaMKII-dependent exocytosis and influx, and decreases through PP1-dependent endocytosis. Free AMPAR on the membrane can bind to scaffolding proteins such as PSD95 to become bound in the PSD region. As mentioned above, the exact dynamics of CaMKII and phosphatase cascades in dendritic spines remain unknown. How those CaMKII and phosphastase dynamics in turn then affect AMPAR dynamics is also unclear.

The various species and their reactions can be found in Tables 1, 2, and 3 for the bistable model, and Tables 4, 5, and 6 for the monostable model.

### Variations of different parameters

To investigate the role of upstream kinase and phosphatase cascades on AMPAR dynamics, we compare two different signaling networks – the bistable and monostable models described above.

Additionally, due to the coupling of CaMKII and PP1 and their opposing effects on AMPAR, we also varied the initial conditions of CaMKII and PP1, specifically their inactive species’ initial condition. This initial concentration variation is meant to represent variations in spine size along with different cytosolic conditions such as variations that occur after synaptic activation or in different regions such as the PSD. Next, we vary the relative contributions of endocytosis/exocytosis and AMPAR influx to investigate the role of signaling on different trafficking mechanisms. This variation of trafficking also accounts for differences in dendritic spine geometry, such as differences in spine surface area and volume that would influence speed or magnitude of the trafficking mechanism.

We also specifically vary the contributions of endocytosis and exocytosis separately in two different manners - 1. where we scale the contributions of the activity dependent terms, and 2. where we scale the contributions of the whole endocytosis and exocytosis terms. These trafficking variations can represent disease states where AMPAR influx, or endocytosis and exocytosis are hindered or upregulated. Finally, we vary the model stimulus by supplying active CaM pulses of different frequencies and magnitudes.

### Numerical methods

The system of deterministic ordinary differential equations was coded and solved in MATLAB R2018b and R2020b. The solver ode23s was used for all numerical simulations. The model was run for 500 s with outputs every 0.01 s. Time stepping was set up automatically in MATLAB. Time-to-peak (TTP) and peak values were found using the MATLAB function max() for a precise numerical value. Steady state was determined when the rate of change of the species was determined to be small (approximately 1 × 10^-3^) or for simplicity at the end of the simulation time range, see Supplemental Material for more information. A sensitivity analysis was run for both signaling models in COPASI. The model files can be found on GitHub under RangamaniLabUCSD.

## Results

Using a compartmental ODE model, we investigated the timescales of different biochemical signaling events underlying AMPAR dynamics during synaptic plasticity. We use the following metrics to generate quantitative comparisons between the different conditions tested in the model: peak concentration, time to peak, steady state concentration, and time to steady state. Bound AMPAR (membrane AMPAR bound to PSD95) is the ultimate model readout for synaptic plasticity.

### Ca^2+^ influx dynamics are the same for both the monostable and the bistable models

We first focus on the early events in the spine in response to Ca^2+^ influx (Figure 2; bistable in blue, monostable in red). When Ca^2+^ floods into the spine, it is rapidly bound by CaM and effluxes through various pumps; see Module 0 and 1 in Table 1 and Table 4. In response to the voltage stimulus (Figure 2a, inset; see Key Components of model), Ca^2+^ influx is rapid and decays within 10 s (Figure 2a), consistent with previous models [10, 14, 21] and experiments [66, 67]. Ca^2+^ bound CaM, denoted as Ca^2+^ CaM, shows a slightly slower increase to peak value compared to Ca^2+^ (290 ms for Ca^2+^CaM versus 90 ms for Ca^2+^) and also decays within the approximately 10 s timescale (Figure 2b). Free CaM can also bind to the scaffolding protein neurogranin [21, 68]. Our model is a closed system for all species except A_mem_ and begins with no CaM·Ng, so this model only considers Ng as a CaM sink. In our simulations, the CaM·Ng complex attains a high value at steady state (9.8 μM) in that same time frame of 10 s (Figure 2c).

Because of the upstream nature of these modules, both the bistable and monostable model show very similar dynamics and model readouts for these species, see Figure 2d-e. We note that the CaM·Ng complex reaches an ultimate peak value after achieving steady state because we define steady state as when the rate of change of the species is small (see Supplemental material). We also see that the time to peak for CaM·Ng in the monostable model is longer than the time to peak for the bistable model (~ 37 s vs ~ 27 s); however the concentrations for peak value and steady state value for both models are all the same (9.81 μM) so the differences in time to peak 2+ and steady state time are not of significance in the model (Figure 2d-e). Thus, the influx of Ca^2+^ and buffering by CaM are not affected by the choice of the model for CaMKII/AMPAR dynamics as expected, establishing the proof-of-principle that the early time scale dynamics, particularly for Ca^2+^ influx, are unaffected by model choice.

### Kinase and phosphatase dynamics differ between the monostable and the bistable models as expected

We next describe the dynamics of the kinases and phosphatases in the cascade for one particular choice of initial conditions (IC) (inactive CaMKII IC of 20 μM and inactive PP1 IC of 0.25 μM; see Table 3 and Table 6). Given the differences in the models used to describe CaMKII activation and phosphatase dynamics, we see substantial differences in the kinetics when comparing the two signaling models (Figure 2). In both models, Ca^2+^ CaM activates CaMKII rapidly to its maximum value (20 μM). However, in the bistable model, CaMKII remains elevated at its maximum value while in the monostable model CaMKII decays towards zero, see Figure 2f. In the bistable model, Ca^2+^CaM activates CaN as the first step of the phosphatase cascade, see Figure 2g-i, blue line. The bistable phosphatase cascade shows rapid activation of CaN to its maximum concentration (1 μM), which then triggers activation of I1 and then PP1 (Figure 2g-i). I1 and PP1 have larger peak times than CaN in the bistable model because of the temporal relay of information (~ 40 s versus less than 1 s for CaN peak), see Figure 2l,m. The monostable model has a different activation pathway for PP1; PP1 is directly activated by Ca^2+^CaM such that it operates on the same timescale as CaMKII [12]. As a result, PP1 and CaMKII have very similar kinetics in the monostable model with a rapid rise followed by a decay to zero (Figure 2f, i, red line).

We compare our computational model results to other experimental [40, 41] and computational results [12] (Figure 2j-k). We note that our goal is not to match the exact dynamics of CaMKII or PP1 dynamics from experiments but to provide estimates for different timescales based on experimental conditions. [41] studies CaMKII dynamics in dendritic spines using fast-framing two-photon fluorescence lifetime imaging of hippocampal slices. Similarly [40] studies CaMKII and various mutants in hippocampal slices using two photon glutamate uncaging. To compare timescales, we scale all other results to our maximum CaMKII and PP1 concentrations. We see that across all published data CaMKII concentration increases rapidly, but there is variation in decay time (17.6 s for [41], 38.1 s for [40], 61.5 s for [12]). The monostable model decays with a rate of 20 s which resembles the rate of [41], while the various other CaMKII results all decay within approximately 100 s except the bistable model which remains elevated (Figure 2j), for this choice of IC. Phosphatase temporal dynamics are harder to compare against experiments because many studies do not capture these dynamics; however PP1 decay parameters appear slower than CaMKII decay [12, 28] so we include these dynamics in our monostable model. Observations of PP1 temporal dynamics from [12] show a longer elevated period compared to our monostable results (decay time of 85.7 s from [12]) but have rise dynamics intermediate between our bistable and monostable dynamics (Figure 2k).

To more directly compare our CaMKII and PP1 temporal dynamics, we compare the decay times across experimental and model results. CaMKII decay dynamics have been estimated experimentally as having two timescales of delay *τ_fast_* = 6.4 ± 0.7 s and *τ_slow_* = 92.6 ± 50.7 s with different fractions of fast and slow decay (74% vs 26%) [41] ([40] shows similar decay values). Equivalent parameters for PP1 could not be immediately found for this system. Our time constants for CaMKII and PP1 decay were *τ* = 20 s and *τ* = 40 s, respectively (See Table 2 and Table 5).

A direct comparison of model readouts for the bistable and the monostable model are shown in Figure 2l-m, respectively. The main differences in the CaMKII dynamics are in the steady state values in the bistable and the monostable model (19.9 μM versus 0.02 μM respectively) and the time to steady state (~ 219 s versus ~ 141 s), where the bistable model takes longer to reach steady state but hovers near maximal value. When comparing PP1 dynamics for both the models, we note that the bistable model has a lower peak value (0.06 μM) when compared to the monostable model (0.25 μM). While PP1 steady state goes to zero for both models for this choice of initial conditions, the time it takes for steady state is different (~ 260 s for the bistable model compared to ~172 s for the monostable model) suggesting that these differences in timescales may have an impact on downstream events.

### AMPAR dynamics are dependent on model choice for CaMKII/PP1 balance

We next investigated how the differences in kinase and phosphatase dynamics might influence the dynamics of AMPAR for a single set of initial conditions (20 μM for CaMKII and 0.25 μM for PP1). Given that downstream signaling can integrate the details of upstream signaling kinetics [69] and can integrate out the effect of upstream timescales [58], we wondered if the differences in the two models would make a difference in AMPAR dynamics. We track three populations of AMPAR in our model – AMPAR in endosomes (Aint, modeled as a volume component), free AMPAR on the synaptic membrane (Amem), and AMPAR bound to PSD95 (bound AMPAR). Internalized AMPAR can become membrane-bound as a result of CaMKII-dependent exocytosis; membrane bound AMPAR is endocytosed in a PP1-dependent manner (R13 in Table 1 and R7 in Table 4). Membrane AMPAR increases through a CaMKII-dependent influx that represents AMPAR diffusing laterally into the spine from extrasynaptic pools of membrane AMPAR outside of the spine [45, 48] (R14 in Table 1 and R8 in Table 4). Membrane bound AMPAR can be immobilized by binding to PSD95 among other PSD proteins (R15 in Table 1 and R9 in Table 4). AMPAR is known to interact with a plethora of cytosolic and membrane bound species at the PSD, often with com-plex phosphorylation or other activation dynamics [49, 51, 70]; however in our system we chose to model PSD95 and Amem binding through simple mass action kinetics (Module 6 in Table 1 and Table 4). Despite simple reaction kinetics, this scaffold interaction significantly influences the system, demonstrating how simple signaling components can dominate more complex signaling motifs [71]. Our simulations show that internal AMPAR (Aint) is quickly exocytosed but then gradually increases again due to endocytosis, with the monostable model having a larger increase in internal AMPAR steady state compared to the bistable model (0.20 μM versus 0.08 μM; Figure 3a). Free AMPAR on the membrane increases to a new steady state with the bistable model having a significantly higher density than the monostable model (Figure 3b). The higher Amem density in the bistable model is because the bistable model has a higher steady state value of CaMKII for the ICs (20 μM for CaMKII and 0.25 μM for PP1) than the monostable model and even though the PP1 peak is higher in the monostable model, the impact of CaMKII is stronger on the balance between Aint and Amem. Naturally, the unbounded scaffolding protein PSD95 decreases to a new steady state value as it binds to free membrane AMPAR (Amem) to form bound AMPAR (Figure S4a) in both cases. We note that membrane AMPAR has rapid binding to form bound AMPAR, but also has a rapid increase in endocytosis/exocytosis and influx rates (Figure S4b-c). Bound AMPAR increases in both models to new steady states but reaches a much higher density in the bistable model compared to the monostable model (~ 1802 receptors per micron squared versus ~ 1229 receptors per micron squared; Figure 3c), again because of the stronger effect of CaMKII. These densities of bound AMPAR are within the range of AMPAR densities observed in experiments [52, 72–77]. The monostable model achieves steady state for all AMPAR species and PSD95 within ~ 117s, faster than the bistable model whose fastest component, Aint, took ~ 125 s to reach steady state (Figure 3d-e).

**Figure 3:**
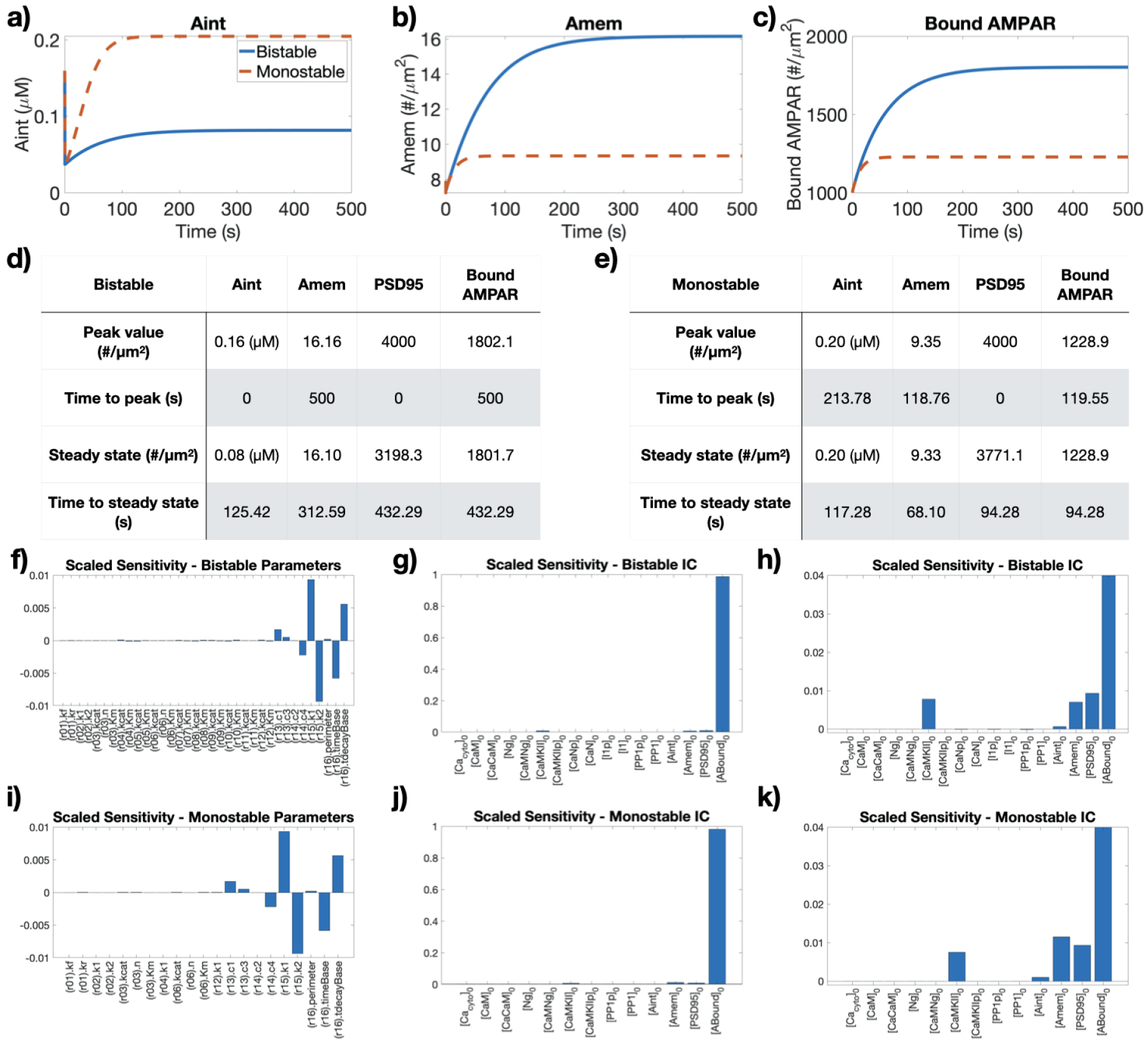
a) Temporal dynamics of internal AMPAR in endosomes (Aint, a), free AMPAR on the membrane (Amem, b), and bound AMPAR (c) for both the bistable and monostable models show similar general trends. Peak and steady state values and times for Aint, Amem, PSD95, and bound AMPAR for the bistable (d) and monostable (e) models show key differences in dynamics. Scaled sensitivity analysis results with respect to the bound AMPAR readout for the bistable parameters (f), bistable initial conditions (g), and a zoomed view of the bistable IC results (h). Scaled sensitivity analysis results with respect to the bound AMPAR readout for the monostable parameters (i), monostable initial conditions (j), and a zoomed view of the monostable IC results (k).

Next, we conducted a sensitivity analysis to better understand the significant parameters and initial conditions that influenced bound AMPAR in both models (Figure 3f-k). The sensitivity analysis was run in COPASI [78] and the influence of each parameter and initial condition was considered on the final readout of bound AMPAR, see Supplemental material sensitivity analysis section for more details. We find similar trends for both the bistable and monostable models – bound AMPAR is most sensitive to the kinetic parameters that govern endo/exocytosis, CaMKII-mediated AMPAR influx, PSD95 binding rates, and to the initial conditions of CaMKII, PSD95, and the subpopulations of different AMPAR species. As expected, the bound AMPAR readout was highly sensitive to the IC of bound AMPAR, but was also sensitive to parameters and species associated with the scaffold binding reactions. Thus, CaMKII-mediated AMPAR influx and AMPAR binding to PSD95 both greatly influence bound AMPAR dynamics as compared to CaMKII and PP1-mediated exocytosis and endocytosis. The slower timescale and higher steady state density for bound AMPAR associated with the bistable model can be explained by the elevated CaMKII steady state that leads to a higher influx of membrane AMPAR. Thus, because of the nature of the upstream kinase (prolonged activation versus finite time activation), the density of bound AMPAR on the membrane can be significantly impacted.

### Effect of CaMKII and PP1 initial conditions on model-dependent AMPAR readout

One of the key differences between a bistable model formulation for CaMKII (in this case coupled with a phosphatase cascade) and the monostable model is the dependence of steady state values for CaMKII and PP1 on the initial conditions (IC) [28, 32]. To understand how bound AMPAR depends on the initial condition of inactive CaMKII and PP1, we conducted a parameter sweep (Figure 4) and focus on the steady state behavior. For temporal dynamics of all active PP1 and active CaMKII for both the bistable and monostable models across the explored IC regimes, see Figure S1. Temporal dynamics of bound AMPAR for both the bistable and monostable models can be found in Figure S3. For the bistable model, active CaMKII, active PP1, and bound AMPAR steady states all show dependence on both CaMKII and PP1 initial conditions, with the higher CaMKII IC leading to higher bound AMPAR density, and higher PP1 IC leading to lower bound AMPAR density (Figure 4a). It is useful to recall that an increase in bound AMPAR density is associated with LTP, while a decrease is associated with LTD. In these cases, despite PP1 IC having a strong influence on the bistable system, all cases either display LTP or no change (NC). High CaMKII IC and low PP1 IC leads to high active CaMKII at steady state in the bistable model and therefore high bound AMPAR, whereas low CaMKII IC with high PP1 IC will lead to low active CaMKII at steady state and therefore low bound AMPAR (Figure 4b). We also see that for two different PP1 concentrations, CaMKII can achieve two different steady state values for the same CaMKII IC, the hallmark of bistability, Figure 4b. Similarly, for two different CaMKII ICs, PP1 can also achieve two different steady state values for the same PP1 IC. Considering bound AMPAR across CaMKII ICs for two different PP1 ICs, for a small PP1 IC (0.1 μM), bound AMPAR at steady state is linear with respect to CaMKII IC; for the higher PP1 IC (0.5 μM), bound AMPAR shows a non-linear dependence on CaMKII ICs. When we vary PP1 ICs for two different CaMKII ICs, we observe that for a small CaMKII IC (12 μM), bound AMPAR steady state decreases with respect to PP1 IC; for the higher CaMKII IC (20 μM), bound AMPAR shows only slight dependence on PP1 IC, slightly decreasing for high PP1 IC.

**Figure 4:**
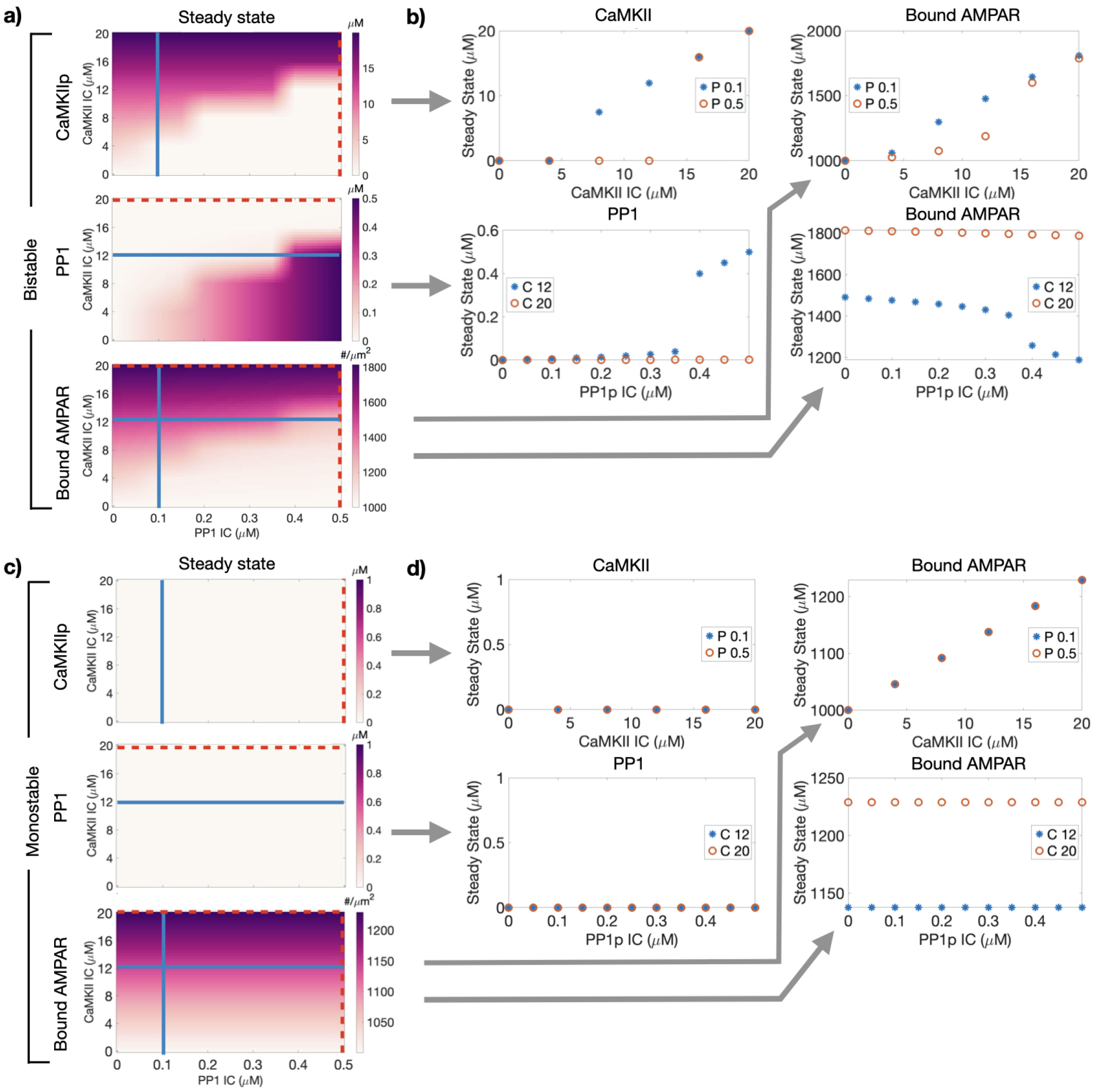
Variations in initial conditions for inactive CaMKII and PP1 influences active CaMKII, active PP1, and bound AMPAR dynamics in a model-dependent manner. a) Steady state values for active CaMKII, active PP1, and bound AMPAR (top to bottom row, respectively) for the bistable model. b) Steady state value of CaMKII, PP1, and bound AMPAR for two set PP1 ICs and varied CaMKII IC and for two set CaMKII IC and varied PP1 ICs for the bistable model. c) Steady state values for active CaMKII, active PP1, and bound AMPAR (top to bottom row, respectively) for the monostable model. d) Steady state value of CaMKII, PP1, and bound AMPAR for two set PP1 ICs and varied CaMKII IC and for two set CaMKII IC and varied PP1 ICs for the monostable model.

We conducted a similar parameter sweep with the monostable model (Figure 4c-d). The monostable model only has one steady state for both CaMKII and PP1 which is zero (Figure 4c). Therefore, the initial condition only determines the peak value of the model without any coupling between CaMKII and PP1, as seen in Figure S2. Bound AMPAR shows dependence on CaMKII IC but none on PP1 IC in this case (Figure 4c,d). Overall the monostable model did not demonstrate a strong dependence on ICs as compared to the bistable model. The steady state heatmaps clearly show that the monostable model has no coupling between CaMKII and PP1 dynamics and the bound AMPAR steady state depends on the CaMKII IC (Figure 4c, d, right column).

### Effect of varying trafficking conditions on AMPAR dynamics

Thus far, we have focused only on the signaling dynamics of CaMKII and PP1 on AMPAR. Next, we investigated how the coupling between signaling dynamics and trafficking dynamics can affect bound AMPAR. As described in Table 1 and Table 4, bound AMPAR also depends on endo- and exocytosis and influx of AMPAR from the extrasynaptic pool (see Figure S4).

The dynamics of *A_mem_* are given by the following equation

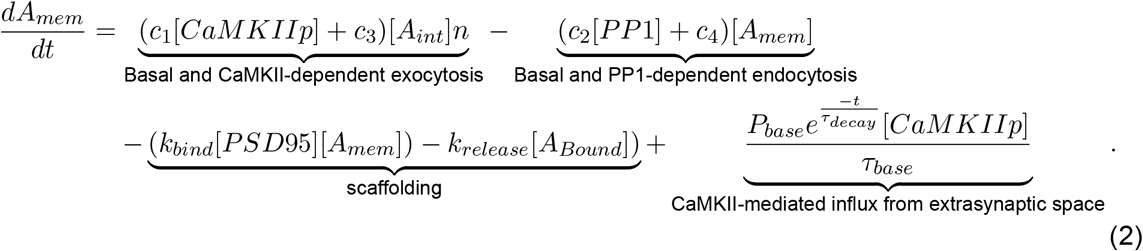

The new terms introduced here are the following: *P_base_* = 0.9069 *μm*, *τ_base_* = 800 s, *τ_decay_* = 60 s, and n = 0.1011 *μm. P_base_* is the spine neck base perimeter and we approximate it from an average thin spine in [10]. Endocytosis and exocytosis rates were taken from [3]. *τ_base_* is the timescale of influx and can be though of as the timescale at which AMPAR moves into the dendritic spine [8]. *τ_decay_* is the timescale of CaMKII activity and captures the timescale over which active CaMKII triggers the influx of membrane AMPAR into the spine [45]. *n* is the lengthscale conversion factor and captures the spine volume to surface area ratio. The CaMKII-mediated influx is inspired by [45] and captures how elevated CaMKII levels can trigger the movement of AMPAR into the activated spine, but the signal must turn off to prevent excessive AMPAR influx. We note that mathematically the presence of the influx term results in an open system for AMPAR mass conservation since the total number of receptors in the simulation can change because of external sources.

The dynamics of bound AMPAR are given by

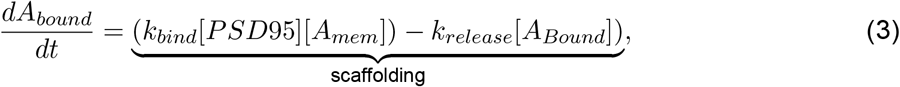

which describes the binding and unbinding of free AMPAR and PSD95, a scaffolding molecule, to form bound AMPAR. We assume that there is a pool of free PSD95 ready to bind membrane AMPAR.

Using this framework, we asked how the different trafficking conditions impact bound AMPAR. In what follows, we compare all the parameter variations to a control, which is defined as the simulation in which all forms of trafficking (PP1-dependent endocytosis, CaMKII-dependent exocytosis, and a CaMKII-dependent influx source of AMPAR) are present (Figure 4a, bottom row for bistable and (Figure 4c, bottom row for monostable). Recall that, for the control cases, bound AMPAR reached a maximum of 1802 per square micron for the bistable case versus 1229 per square micron for the monostable case.

We considered the steady state value (value at 500 s) of all AMPAR species for two *in silico* knockout trafficking conditions - no CaMKII-mediated influx (Figure 5a) and no endocytosis/exocytosis (Figure 5b) for different CaMKII and PP1 ICs. Temporal dynamics of bound AMPAR with different CaMKII and PP1 initial conditions for the various trafficking knockout cases can be found in Figure S3 and heatmaps showing PSD95 steady state dynamics for different CaMKII and PP1 initial conditions for the various trafficking cases are in Figure S5.

**Figure 5:**
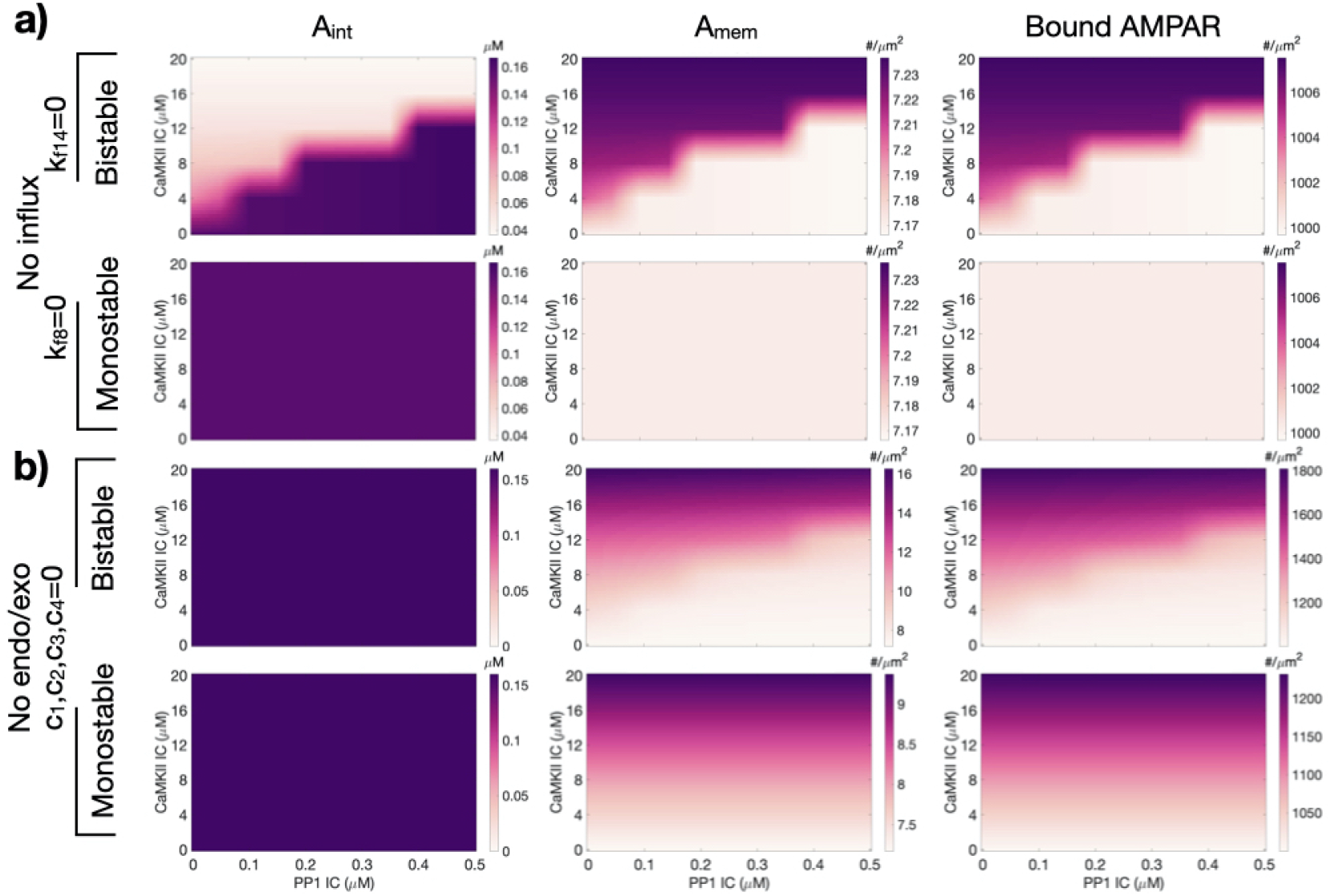
Effect of CaMKII and PP1 initial condition on trafficking mechanisms. a) Species value at 500 s for internal AMPAR (Aint), membrane AMPAR (Amem), and bound AMPAR for the bistable (top row) and monostable (bottom row) models in the trafficking case without an external AMPAR source for different CaMKII and PP1 ICs. b) Species value at 500 s for the bistable (top row) and monostable (bottom row) models in the trafficking case without endocytosis and exocytosis for different CaMKII and PP1 ICs.

We first asked what happens if influx is removed as a source of AMPAR but endocytosis and exocytosis are retained? We found that the contribution of endocytosis and exocytosis were smaller compared to the influx from extrasynaptic regions (Figure 5a). The bistable model shows dependence on both CaMKII/PP1 IC, while the monostable case leads to homogeneous results for all species (Figure 5a). Thus, in the absence of influx but in the presence of endocytosis and exocytosis, we find that bound AMPAR readout mimics the upstream signaling pathway. The steady state for both Amem and bound AMPAR was lower in the monostable case versus the bistable case, highlighting how the elevated CaMKII levels in the bistable case contribute to both free and bound AMPAR pools even in the absence of endo and exocytosis. However, if we retain influx but turn off endocytosis/exocytosis, internal AMPAR cannot be trafficked onto the membrane, so there is no change in Aint concentration for either choice of upstream signaling (Figure 5b, first column). Additionally, there is a coupled dependence on IC for Amem and bound AMPAR for the bistable case but only CaMKII IC dependence for Amem and bound AMPAR for the monostable case (Figure 5b). Because there was no direct impact of PP1 and CaMKII on Amem levels through endocytosis and exocytosis, the coupled PP1/CaMKII effect in the bistable model is due to the upstream interactions between CaMKII and PP1 that then influence steady state CaMKII levels that mediate AMPAR influx. In the parameter space of varying CaMKII and PP1 ICs, in the bistable model, we observe a smooth dependence of Amem and bound AMPAR; this is different from the case without influx but with endo/exocytosis where the bound AMPAR demonstrate a step-like response. This suggests that influx of AMPAR term takes a bistable cascade as an input and produces a proportional mapping as an output.

Is it possible that the conclusion that influx of AMPAR is a dominant contributor to bound AMPAR dynamics is a result of the choice of parameters for the endo/exocytosis terms (obtained from [30])? And can we investigate the endocytosis and exocytosis contributions separately as suggested by [79]? Recall that the exocytosis and endocytosis term is given as

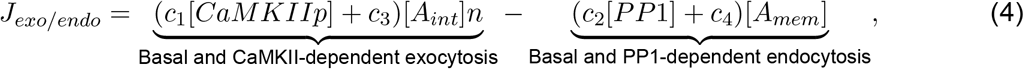

where both exocytosis and endocytosis have a species dependent component and a basal contribution. To answer the questions raised above, we varied the relative contributions of CaMKII-mediated exocytosis and PP1-mediated endocytosis in two different ways. In the first method, we scale the c_1_ and c_2_ terms, such that

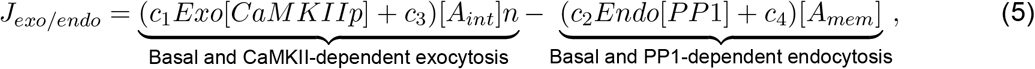

where Exo and Endo are scaling factors that range from 0 to 500 and essentially modify the rate of activity-dependent endocytosis or exocytosis and can be attributed to other biophysical factors that affect trafficking [57, 80]. A maximum value of 500 was chosen to ensure that given the control initial conditions of CaMKII and PP1, the endocytosis term could achieve the same magnitude as the original exocytosis term (Figure 6a and c). In this case, the stimulus was applied at t = 0, since the system was already at steady state.

**Figure 6:**
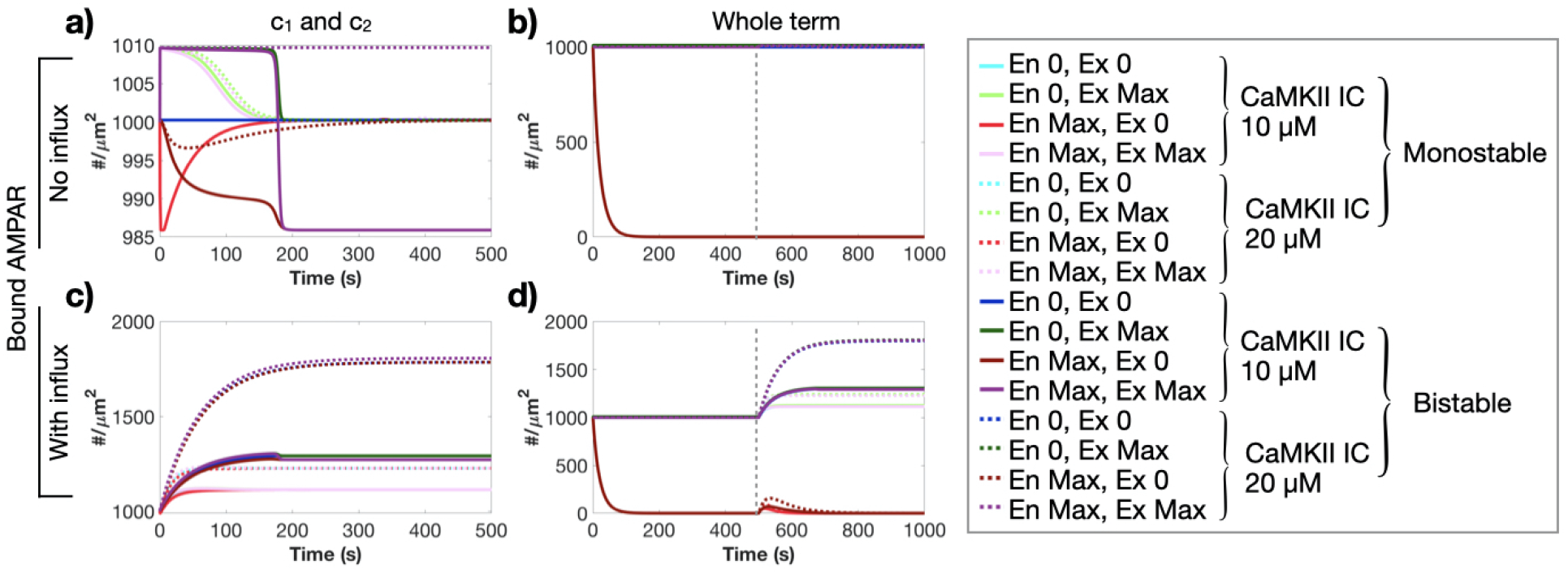
Variation of different trafficking conditions on bound AMPAR. Temporal dynamics of bound AMPAR without AMPAR influx for two different CaMKII initial conditions (10 μM and 20 μM), different contributions of endocytosis and exocytosis, for the bistable and monostable models, for the scaling method that scales c_1_ and c_2_ (a) and the whole endocytosis and exocytosis terms (b). Temporal dynamics of bound AMPAR with AMPAR influx for two different CaMKII initial conditions (10 μM and 20 μM), different contributions of endocytosis and exocytosis, for the bistable and monostable models, for the scaling method that scales c_1_ and c_2_ (c) and the whole endocytosis and exocytosis terms (d). Endocytosis and exocytosis were varied in two different ways: a,c) (c_1_ and c_2_): the species dependent rates of c_1_ and c_2_ were scaled; and b,d) (Whole term): the entire exocytosis and endocytosis term was scaled (c_1_, c_3_ and c_2_, c_4_). The dashed gray line in b and d indicates when the stimulus was applied to the system at t = 500 s. Gray inset: Legend where general color corresponds to a predetermined endocytosis (En)/exocytosis (Ex) scaling in this order (En 0, Ex 0; En 0, Ex Max; En Max, Ex 0; En Max, Ex Max), solid and dashed lines indicate the 10 μM and 20 μM CaMKII IC, respectively; and the lighter colors indicate the monostable model and dark colors indicate the bistable model. Note Max is 500 for a and c, and 1 for b and d.

In the second method, we scale the entire exocytosis and endocytosis terms, such that

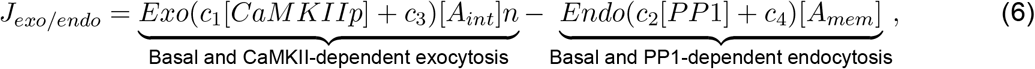

where Exo and Endo are scaling factors that range from 0 to 1 and modify the cumulative rate of endocytosis or exocytosis. This scaling essentially alters the steady state of the system in the absence of stimulus and so we first simulated the system to steady state and then added the stimulus at t = 500 s (gray dashed line in Figure 6b and d).

We considered four different cases for both the bistable and monostable models, Figure 6. First we considered the models with and without influx (Figure 6, top and bottom rows, respectively). Second we considered two different initial conditions for CaMKII, 10 and 20 μM, but only one IC for PP1 (0.25 μM). Recall that for the bistable model, the first condition (10 μM CaMKII and 0.25μM PP1) leads to an elevated PP1 concentration, while the second condition (20 μM CaMKII and 0.25 μM PP1) leads to an elevated CaMKII concentration. We plot bound AMPAR over time for four different endocytosis (En) and exocytosis (Ex) scaling sets (En 0, Ex 0; En 0, Ex Max; En Max, Ex 0; En Max, Ex Max) for both scaling methods, Figure 6. For temporal plots of bound AMPAR for each of these various conditions, see Figure S8. For a closer analysis of AMPAR dynamics for the c_1_ and c_2_ approach, see Figure S7. For steady state values of bound AMPAR for both approaches, see Figure S6.

In the absence of influx of AMPAR, we find that bound AMPAR can achieve a new steady state after stimulus if there is a constant elevated concentration for CaMKII or PP1 that drives either endocytosis or exocytosis, Figure 6a,b. The monostable case does not have sustained CaMKII and PP1 activation (Figure 2f,i) and therefore, in all variations the monostable model returns bound AMPAR to its initial condition. In contrast, the bistable model has elevated PP1 for the 10 μM CaMKII IC, and elevated CaMKII for the 20 μM CaMKII IC; depending on the dominant contribution between active endocytosis and exocytosis, bound AMPAR can attain a new increased or decreased steady state. When the entire terms are scaled, for the case when the relative contribution of endocytosis is maximized for both values of CaMKII IC, bound AMPAR steady state even before stimulus goes to zero (Figure 6b; red lines). This suggests that in the absence of influx, even for situations where CaMKII is active at high concentrations at steady state, it is possible to get low bound AMPAR because of the contributions of trafficking.

When AMPAR influx is present, bound AMPAR demonstrates a dose-dependence on CaMKII as opposed to a switch-like response, Figure 6c,d. When the entire trafficking terms are scaled and endocytosis is maximized for either CaMKII IC, we see that with an influx the system has a transient increase in bound AMPAR after stimulation at t = 500 s before it returns back to its previous zero steady state (Figure 6d, red lines). Thus, we find that even when influx is present, a balance between endocytosis and exocytosis is necessary to elevate bound AMPAR on the membrane.

### Bound AMPAR as a function of time-dependent stimuli

Thus far, we have focused on model choices and variations for a single stimulus, see Figure 2a, inset. In reality, dendritic spines receive a sequence of stimuli in a frequency-dependent manner [12, 27, 58]. Here we consider the effect of a train of elevated Ca^2+^ CaM activity at different spiking frequencies (0.5 Hz, 0.1 Hz, 0.05 Hz, [81]), for two different stimulus strengths and two different CaMKII ICs, see Figure 7a. Previous studies have shown that CaMKII can integrate calcium/calmodulin activity as a leaky integrator [21, 41], so we consider how stimulus magnitude and frequency affect CaMKII and bound AMPAR dynamics. We note that for the bistable model, there was no noticeable difference in CaMKII and bound AMPAR dynamics for the different combinations of conditions, see Figure S9. This is because the activation of CaMKII and bound AMPAR is already maxed out due to the first activation spike so subsequent spikes have no effect. In the case of the monostable model, we observe that CaMKII behaves as a leaky integrator as the stimulus frequency increases (Figure 7b) and the peak values are proportional to the CaMKII IC. Bound AMPAR correspondingly shows a frequency and CaMKII IC-dependent response (Figure 7c). When we look at the steady state values of bound AMPAR, we observe that for 1X stimulus, the value of bound AMPAR decreases as the frequency increases for both values of CaMKII IC (Figure 7d). When we increase the stimulus amplitude by 10-fold (blue line in Figure 7a), the monostable model shows similar temporal dependencies for CaMKII (Figure 7e) with the peak values maxing out the available CaMKII. Bound AMPAR integrates these temporal dynamics and shows higher values than the 1x stimulus (Figure 7f). The value of bound AMPAR, in this case of 10X stimulus, increases as the stimulus frequency increases for both values of CaMKII IC (Figure 7g).

**Figure 7:**
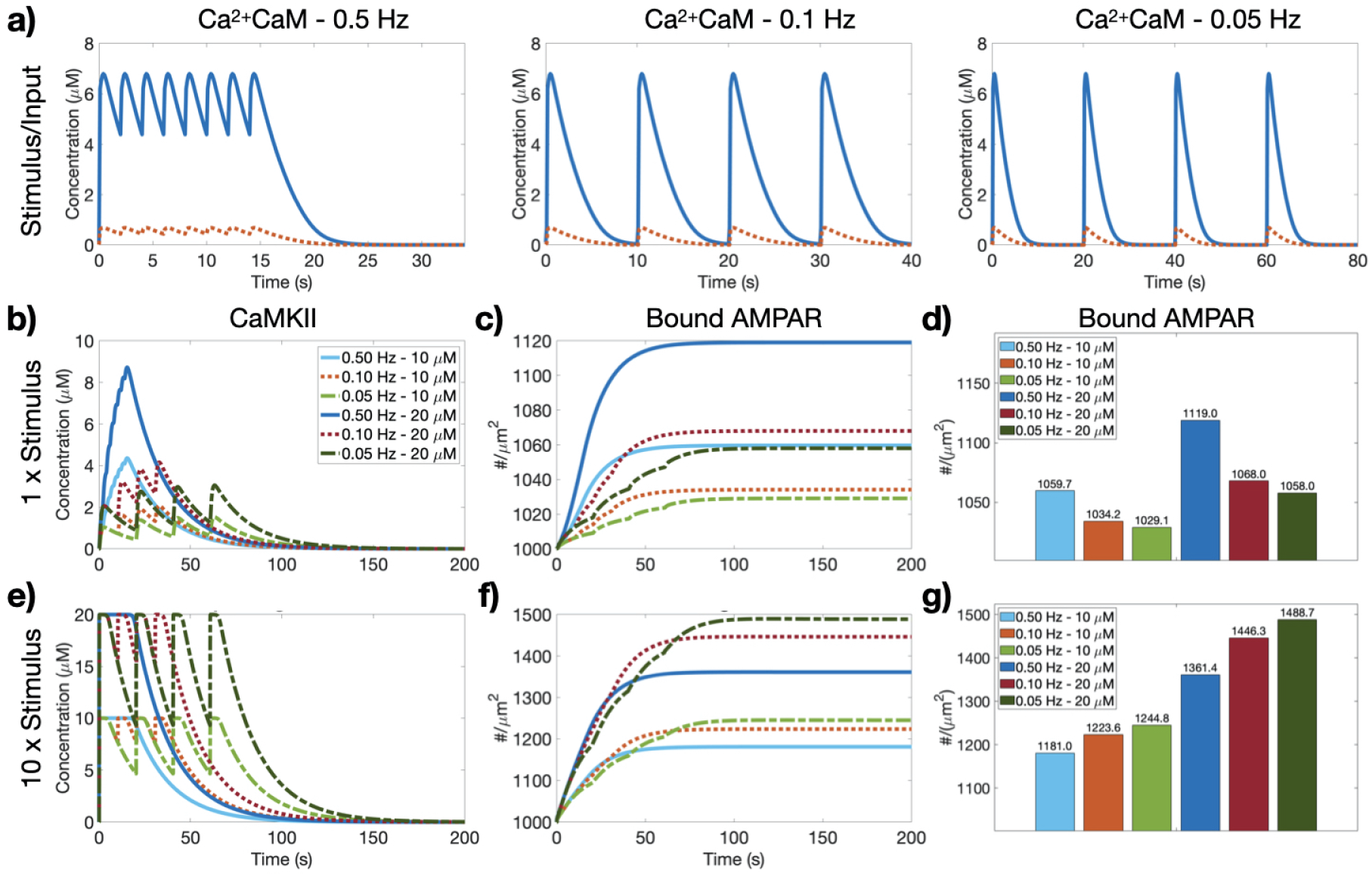
Effect of multiple active Calmodulin spikes on CaMKII and bound AMPAR dynamics in the monostable model. a) Multiple spikes of active CaM were input into the different model systems at different frequencies (0.5 Hz, 0.1 Hz, 0.05 Hz; left, middle, and right, respectively) at two different amplitudes (large - blue, small - red). The large stimulus is 10 times the smaller stimulus. Active CaMKII (b) and bound AMPAR (c) temporal dynamics for the monostable model for the small Ca^2+^CaM stimulus for all three frequencies and both CaMKII IC. d) Steady state bound AMPAR for the small stimulus for each frequency and CaMKII IC. Active CaMKII (e) and bound AMPAR (f) temporal dynamics for the monostable model for the large Ca^2+^CaM stimulus for all three frequencies and both CaMKII IC. g) Steady state bound AMPAR for the large stimulus for each frequency and CaMKII IC.

## Discussion

The choice of model formulation is critical for the interpretation of results in computational and systems biophysics. In the case of synaptic plasticity and learning models, the idea that CaMKII can act as a molecular marker for learning and that a bistable model could be interpreted as a molecular switch has received a lot of attention [59]. Indeed, the idea of a bistable switch has the appeal of making direct associations with LTP and LTD – high stable CaMKII can be interpreted as LTP and low stable CaMKII can be interpreted as LTD [22]. There are two scenarios that complicate this simple picture. First, detailed kinetic models of CaMKII monomers and holoenzymes and experiments have not shown evidence of CaMKII bistability [21, 26, 33]. Second, in recent years, it has become clearer that the increase of AMPAR density at the PSD is perhaps a better molecular marker for learning [82–85] and the net increase in AMPAR depends on various trafficking modalities which in turn depend on upstream signaling pathways. Therefore, here, we chose to investigate how the model choice for upstream signaling could affect the predictions of the model for bound AMPAR, Figure 8a. Additionally, many neurological disorders and diseases include a component of AMPAR dysregulation, Figure 8b. For example, Huntington’s disease involves a disruption in the binding between AMPAR and scaffolding molecules such as PSD-95; both Alzheimer’s Disease and Parkinson’s Disease related dementia involve an increase in AMPAR endocytosis in response to A*β* oligomers, while Parkinson’s Disease also involves an impairment in scaffold binding and AMPAR endocytosis; and mental stress is believed to cause increases in AMPAR lateral diffusion [86]. Therefore, it is clear that the regulation and dynamics of AMPAR have significant neurological consequences. By considering two different signaling model candidates and performing signaling and trafficking variations Figure 1, we can provide the following predictive insights from our model.

**Figure 8:**
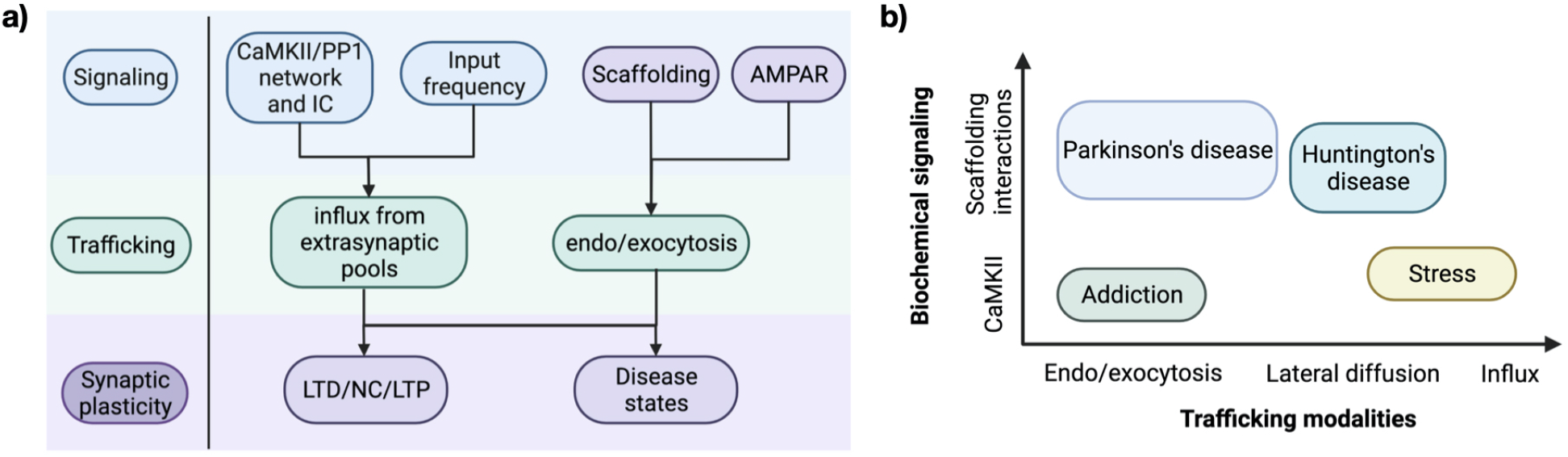
Synaptic consequences of coupling between AMPAR signaling and trafficking. a) Signaling and trafficking components interact to determine changes in synaptic plasticity, whether LTD, NC, LTP, or disease states. b) Dysfunction within either the AMPAR signaling or trafficking mechanisms can lead to various disease states, including Parkinson’s disease, Huntington’s disease, addiction, and stress. Figure made in Biorender.com.

The first insight is that bound AMPAR mimics the dynamics of the upstream model (bistable or monostable) with respect to endo/exocytosis rates as well as ICs in the absence of AMPAR influx. We found that the different signaling models (Figure 2) translated to different AMPAR timescales (Figure 3), primarily through their CaMKII dynamics which informed the influx of AMPAR. Indeed, variations of different parameters and coupling within each model showed that by construction, the bistable model had a stronger coupling between bound AMPAR and upstream signaling than the monostable model at steady state, Figure 4. These pattern held when considering each signaling model with endocytosis and exocytosis or with the AMPAR influx, separately, Figure 5. Thus, careful dissection of the upstream signaling pathways and crosstalk is necessary for building predictive models.

Second, by changing the relative contributions of trafficking, we predict that effects of trafficking can dominate the signaling effects. Specifically, our model predicts that influx of AMPAR from the extrasynaptic space serves to partially override the impact of upstream signaling and trafficking, Figure 6. In some cases but not all, where endocytosis dominates, where one would expect LTD-like behavior, influx of receptors allows for bound AMPAR to either increase or remain unchanged, thereby preventing LTD. Additionally in Figure 6d, when bound AMPAR started at zero but had an AMPAR influx, bound AMPAR transiently increased due to this AMPAR influx but the large endocytosis term drove the system back to zero steady state, (Figure 6d, red lines). This is the only condition that does not show an increase in bound AMPAR steady state when there is an influx term, and could be representative of a disease state where the spine remains silent despite AMPAR influx [79]. Coupling of trafficking with signaling is critical to build mechanochemical models of complex biological processes. Even though we use simple rate representations of trafficking, these rates can be interpreted as factors outside of the signaling pathway such as membrane tension [87–89], lipid composition [90], and other mechanical factors [91, 92]. The third prediction, related to the previous insight, is that in high CaMKII conditions, the system can lose dependence on endo/exocytosis levels because CaMKII can determine the AMPAR outcome through its influence on the influx term or maxing out the exocytosis term, demonstrating how signaling can drive overall dynamics, as seen in the bistable model with high CaMKII IC (see Figure S6a, left column).

The fourth prediction from the models is that for bound AMPAR to achieve a new steady state value, it requires one of two things. Option 1 – seen best through the with influx cases in Figure 6 – is an increase in the total AMPAR in the system (increased membrane AMPAR through a source term, an open system). Option 2 – The presence of a constantly driving term in the endo/exocytosis rates - either elevated PP1 or elevated CaMKII. These can be seen through the difference in bistable and monostable model behavior in Figure 6. The monostable model showed no dependence on endo/exocytosis when there was no influx; instead, it needed an influx of membrane AMPAR to shift the system to a new steady state value. This is why the bistable case can achieve new steady state values despite no influx, while the monostable case cannot.

Finally, we found that the monostable and bistable models had different effects on integrating time-dependent Ca^2+^ influx (Figure 7). For example, the bistable model was not very sensitive to temporal changes to the stimulus (Figure S9). This is because once CaMKII reached a high steady state, it remained at that value. The monostable model on the other hand demonstrated the ability to (a) integrate input frequency and filter out the high frequency effects while retaining sensitivity and (b) show a stimulus-dependent bound AMPAR response. Thus, we predict that while bistable models of CaMKII have appeal in terms of ‘on’ and ‘off’ switches, a monostable model has a distinct advantage of being able to respond proportionally to time-dependent input. This is a desirable trait in synaptic plasticity since frequency dependence is a huge part of learning and memory formation [93]. It has been known that frequency of stimulation can play important roles in CaMKII activation and induction of different synaptic plasticity forms [27, 94]. Historically, many models of synaptic plasticity have relied on the integration of fast variables by slower parameters [13, 18, 58, 94, 95]. Therefore, we found that integration of fast parameters by slow species in synaptic plasticity is actually model dependent. This indicates that there can be more nuance to synaptic plasticity depending on its underlying signaling network dynamics [94]. Therefore, a key takeaway from these results is that the underlying parameters and model architecture can influence not just synaptic plasticity readouts but their dependence on upstream signaling.

Here we used a simplified signaling network to investigate a complex biological process, a method that has successfully explained complex signaling dynamics in dendritic spines before [96]. We acknowledge that our model has many simplifying assumptions and adding biochemical details at various levels is required to build more confidence in these approaches. In addition to biochemical complexity, we will also need to consider other factors that are known to affect synaptic plasticity into the computational models. For instance, signaling nanodomains exist within dendritic spines [97]. Often driven by Ca^2+^ influx, these regions of activation can trigger localized events and it is hypothesized that CaMKII could behave differently in localized domains versus whole dendritic spines [37, 98]. The PSD region also acts as a protein dense, membraneless compartment that can localize signaling processes due to enriched protein concentrations [99]. The effects of this localization and crowding are not clear; for example, while PP1 is enriched in the PSD, it is possible that crowding and additional molecular binding partners can shield CaMKII from dephosphorylation from PP1 in this region [43, 61]. Neurogranin (Ng) plays a seemingly contradictory role in dendritic spines in that despite binding CaM and thus reducing the amount of free CaM available in the system, its presence is known to lower the threshold of LTP through the NMDAR/CaMKII pathway [68, 100]. This contradiction has been explained as follows – Ng-mediated CaM sequestration in the dendritic spine effectively increases the available CaM during spine activation [21, 68]. Specifically Ng can localize CaM close to the plasma membrane of spines to enhance CaMKII activation over CaN activation, leading to enhanced synaptic strength [21, 68, 100]. Additionally, PSD95 itself gets phosphorylated and that affects accumulation at the PSD, but this signaling occurs through a different pathway [51] compared to the signaling considered in this model; therefore, we do not include PSD95 phosphorylation in this study. However, activation and localization of PSD95 and other important scaffolding species could play important roles in AMPAR dynamics at the PSD. Therefore, the localization of these subdomains can play key roles in regulating synaptic plasticity and can actually help explain some of the conflicting experimental evidence on CaMKII dynamics [101].

Trafficking also complicates matters for signaling as both PP1 and CaMKII are recruited into the spine in response to NMDAR activity [23, 43]. While we have considered some trafficking of AMPAR into the synapse through either CaMKII-mediated membrane influx or endocytosis and exocytosis, there is spatial complexity associated with trafficking of AMPAR because of lateral membrane diffusion of AMPAR from the perisynaptic region and from extrasynaptic pools both in the spine and on the dendrite, because of cytosolic influx of endosomes outside of the spine, and because of specific locations of endocytosis/exocytosis [49, 102, 103]. A complication of AMPAR trafficking is that dendritic spine geometry has been found to alter AMPAR diffusion into dendritic spines, making spine geometry another important factor to investigate [8]. An important assumption in this work is that in increase in AMPAR is from AMPAR on the membrane diffusing laterally into the spine or PSD. This analysis would be different if the influx term instead affect internal AMPAR (Aint) which can only become bound AMPAR by exocytosing onto the membrane; or if there was a combination of influx through endosomes in the cytosol and laterally diffusing membrane AMPAR. Thus, spatial models that take these effects into account would be important to fully explore these details.

In summary, we have demonstrated how two different simple biochemical signaling pathways of CaMKII and PP1 can lead to different AMPAR dynamics downstream, Figure 8a. From a signaling perspective, our results highlight the importance of network architecture, the strength of molecular interactions, and the reaction dynamics. With regards to synaptic plasticity, our results highlight the nuances to LTP induction and how modeling efforts of LTP need to be cognizant of the many dependencies of synaptic plasticity (e.g. stimulus magnitude and frequency, upstream signaling dynamics and concentrations, signaling nanodomains, geometric considerations, etc). Both of the CaMKII/PP1 models investigated in this work are possible candidates for what occurs in dendritic spines and from our results, both probably exist as the monostable model acts as a CaMKII leaky integrator [21, 41] but CaMKII is known to autophosphorylate as in the bistable model [25]. The two different signaling dynamics might be differentiated between spatially in nanodomains of activity where conditions favor one versus the other, or they might have different triggers [104]. Nonetheless, these results indicate that the induction and regulation of synaptic plasticity, in this case LTP, might be much more nuanced than previously assumed, and will require a more thorough investigation to parse out the different signaling, spatial, and temporal influences behind synaptic plasticity.

## Acknowledgements

We thank members of the Rangamani Lab for their comments and support. This work was supported by a National Defense Science and Engineering Graduate (NDSEG) Fellowship to M.K.B., and Air Force Office of Scientific Research FA9550-18-1-0051 to P.R.

## Contributions

M.B. and P.R. conceived the project. M.B. conducted simulations. M.B. and P.R. conducted data analysis. M.B. generated all figures. M.B. and P.R. wrote the manuscript.

## Supplemental Material

### Equations for Compartmental Model

We constructed a compartmental ordinary differential equation model of signaling networks describing AMPAR dynamics in response to a single calcium influx event in a dendritic spine of a hippocampal pyramidal neuron. The model is comprised of two compartments - the cytoplasm and the plasma membrane. The dendritic spine plasma membrane surface area was taken to be 0.8 *μm*^2^ which corresponds to an average volume dendritic spine of approximately 0.06 *μm*^3^. The lengthscale conversion between membrane and volume reactions was taken as 0.1011 *μm*, which was the averaged volume to surface area ratio of a realistic dendritic segment of a hippocampal neuron [54]. All reaction equations, parameters, and initial conditions are presented in the tables in the main text (Tables 1,2,3,4,5,6). Initial conditions for the cytosolic species were taken from [10, 28], and initial conditions for the membrane species were approximated from [9, 49, 70, 105]. Various reaction parameters were taken from [3, 10, 12, 28, 41].

### Calcium influx module

Calcium influx is driven by a voltage depolarization due to a excitatory postsynaptic potential (EPSP) and back propagating potential (BPAP) separated by 2 ms. Calcium comes into the spine through NMDAR and VSCC, and is pumped out of the spine through PMCA and NCX pumps. Additionally there is a correcting leak term. Calcium binds to CaM to form Ca^2+^CaM (Ca/CaM complex). The various calcium flux terms are modified to have volumetric reaction units and are taken directly from [10]. See [10] for a sensitivity analysis of the different parameters in the calcium influx module.

### Calmodulin, Kinases, and Phosphatases Modules

Calcium influx leads to activation of downstream signals such as CaM, CaN, CaMKII, and PP1. CaM is activated by binding to multiple Ca^2+^ ions and can be described as a multistage process 2+ where both CaM bound to either 2 or 4 Ca^2+^ ions can lead to downstream signaling activation [21, 27]. However, for simplicity, we approximate the reaction as a single binding event of CaM binding to 3 Ca^2+^ ions [28]. CaM also binds to neurogranin which acts as a CaM sink [106]. CaMKII and the phosphatase cascade have different equations for the bistable [28, 30] versus monostable models [41].

### For the bistable model

The bistable model is termed bistable because the system can produce multiple steady states for either CaMKII and PP1. CaMKII is activated by the Ca^2+^ CaM complex and then can autophosphorylate itself. Active CaMKII is deactivated by active PP1 [19, 28, 30]. CaN is activated by Ca^2+^CaM complex and is deactivated by active CaMKII. Active CaN then activates I1 which is also deactivated by active CaMKII. This I1 species can be thought of as a representation of I1 downstream effects rather than the exact dynamics of I1. I1 inhibits PP1 activity [107], so this species can be thought of as the result of CaN deactivating I1 which then would trigger an increase in PP1 activity [31]. Therefore, this I1 representative species activates PP1. PP1 will then autoactivate itself and be deactivated by active CaMKII [28, 30].

### For the monostable model

The monostable model has Ca^2+^CaM complex activate both CaMKII and PP1 directly. Both species then decay exponentially with linear dependence on their own concentrations [26, 33, 62].

### AMPAR and scaffold module

AMPAR exists in both compartments - Aint in the volume, and Amem and Abound on the membrane. Aint represents AMPAR in endosomes in the cytoplasm. Amem represents free AMPAR on the plasma membrane. AMPAR interacts with many different proteins in the PSD region, but here we specifically model PSD95 on the PSD membrane where it binds membrane AMPAR and localizes it to the PSD region. PSD95 has been found to be vital for AMPAR localization to the PSD [22, 70]. Abound represents membrane AMPAR bound to PSD95.

On the membrane, Amem undergoes endocytosis and exocytosis and has an active CaMKII-dependent boundary flux at the base of the spine neck. The equation for Amem is given by

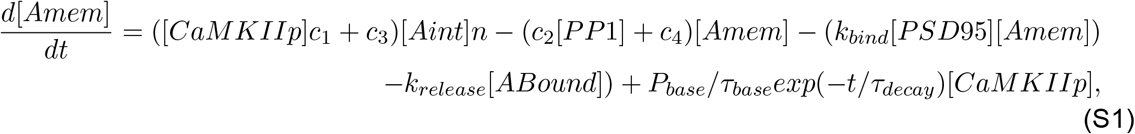

where *P_base_* = 0.9069 *μm, τ_base_* = 800 s, *τ_decay_* = 60 s, and n = 0.1011 μm is the spine neck base perimeter, timescale of influx, timescale of CaMKII activity, and the lengthscale conversion factor, respectively. This CaMKII-mediated influx is inspired by [45]. Endocytosis and exocytosis rates were taken from [3]. Endocytosis and exocytosis affects both Aint and Amem. The reactions for Aint have units of 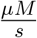, while the reactions for Amem has units of 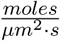. Therefore, we use the lengthscale factor n to convert between units for these reactions.

### Trafficking conditions and model variations

We investigate two forms of AMPAR trafficking during LTP - endo/exocytosis and influx from extrasynaptic pools of membrane AMPAR outside of the spine. We considered the control case when both trafficking modalities were present, the no endo/exocytosis case when there was only a CaMKII-mediated influx, and the no influx case when there was only endo/exocytosis. We also varied the initial condition of inactive CaMKII and inactive PP1 from 0-20 μM and 0-0.5 μM, respectively. We varied contributions to endocytosis and exocytosis from 0-500 when scaling c_1_ and c_2_, and from 0-1 when scaling the whole endocytosis and exocytosis terms. We also varied the frequency of input to the model by applying spikes of Ca^2+^CaM at different frequencies to the model. Therefore, the stimulus of the simulations in Figure 7 and Figure S9 is Ca^2+^CaM, instead of the previous Ca^2+^ influx.

### Numerical details

We ran all model results in MATLAB 2018b or 2020b, and conducted the sensitivity analysis in COPASI. For time to peak and max values of different species, we used the function max() in MATLAB. For Figure 2 and Figure 3, steady state was designated as when the rate of change was below 1 × 10^-3^ for all cytosolic species except Aint and the membrane species. We designate steady state for Aint, Amem, PSD95, and Abound as when the rate is below 1 × 10^-4^, 1 × 10^-3^, 5 × 10^-3^, and 5 × 10^-3^, respectively. If time to steady state and steady state value is not achieved, then we take the value at the end of the simulation peak (500 s). For Figure 4 and Figure 5, steady state is taken as the value at 500 s.

### Sensitivity Analysis

We conducted a sensitivity analysis for both the bistable and monostable models with respect to the final readout, bound AMPAR. The analysis was run in COPASI with respect to all model parameters and initial conditions (Figure 3f-k). Both models showed similar sensitivities to parameters and initial conditions. The most sensitivity was shown toward the PSD95 and AMPAR species initial conditions and parameters, and the AMPAR influx terms. By far the most sensitivity was towards the initial condition for bound AMPAR which is expected. Both models were also sensitive to the reaction rates for membrane AMPAR binding to PSD95 and AMPAR influx. Lastly, the models showed sensitivity to the initial condition of inactive CaMKII, which matches our findings.

## Supplemental Figures

### Variations in CaMKII/PP1 initial conditions alter CaMKII/PP1 and AMPAR dynamics in a model dependent manner

We vary the initial conditions of CaMKII and PP1 for both models to observe the effects on CaMKII, PP1, and bound AMPAR dynamics, Figure S1 and Figure S3. In the bistable model, CaMKII and PP1 can achieve different steady states depending on IC (ex. steady states of 12 vs 20 μM and of 12 vs 0 μM for CaMKII, steady states of 0 versus 0.5 μM for PP1; Figure S2b, left and middle column). Furthermore, for the same CaMKII IC but different PP1 IC, bound AMPAR increases with the same dynamics initially before diverging to a lower steady state for the higher PP1 condition. We quantify the peak values and time to peak values for a range of CaMKII and PP1 initial conditions (0-20 μM and 0-0.5 μM, respectively) for both the bistable and monostable models in Figure S2. For the bistable model, active CaMKII peak value shows no dependence on PP1 IC, while time to peak value shows dependence for low PP1 IC and low CaMKII IC; this is because for the zero CaMKII IC, the peak value is taken as time zero. CaMKII peaks rapidly for all IC for the bistable model (before 500 ms). CaMKII steady state for the bistable model shows dependence on both CaMKII and PP1 IC and ranges from 0 - 20 μM (Figure 4). In the bistable model, PP1 dynamics and steady states are also different based on both the CaMKII and PP1 IC (Figure 4). The PP1 peak value time does show some variations in the CaMKII/PP1 dependence trend with a few regions of slower peak value times for lower PP1 ICs at both low and high CaMKII IC (ex. CaMKII IC 4 μM and PP1 IC 0.05 μM; Figure S2a, middle row and right column). PP1 steady states also range across the whole accessible concentration regime from 0 - 0.5 μM (Figure 4). These different CaMKII and PP1 dynamics translate to different bound AMPAR dynamics in the bistable model for the different IC combinations; in particular, higher CaMKII IC lead to higher bound AMPAR peak values that took longer to reach steady state, due to the increased CaMKII-mediated AMPAR influx. The time to peak value was particularly striking, with a clear distinction between CaMKII dominance and PP1 dominance (diagonal stepping line, Figure S2).

**Figure S1:**
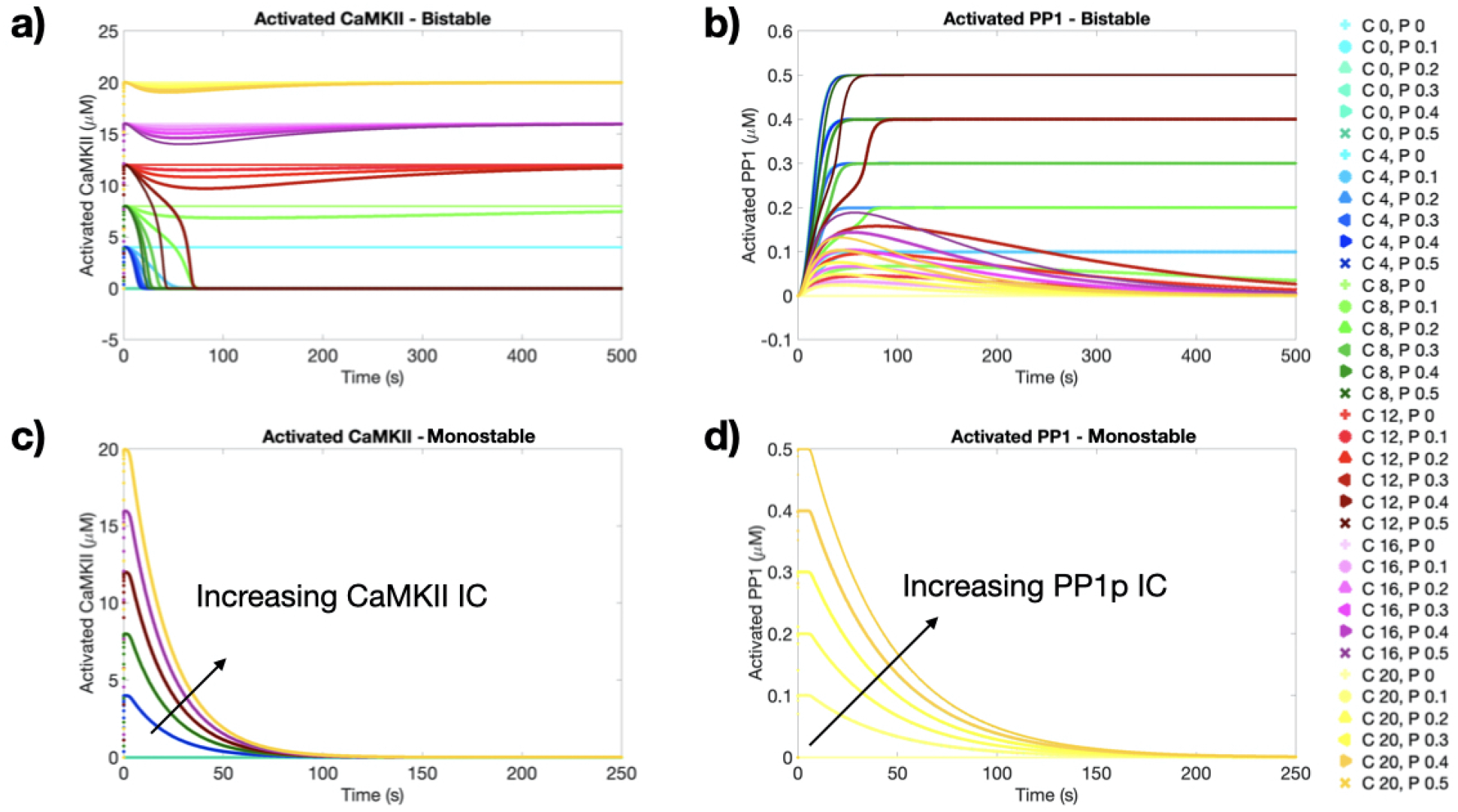
Temporal dynamics for CaMKII and PP1 for both the bistable and monostable models for all CaMKII and PP1 IC combinations. Active CaMKII (a) and active PP1 (b) dynamics for the bistable model show that the coupling between CaMKII and PP1 allows for a variety of different steady states dependent on IC. Active CaMKII (c) and active PP1 (d) dynamics for the monostable model show that CaMKII and PP1 dynamics depend only on their own respective concentration and all species decay back to zero. The C and P in legend stand for CaMKII and PP1, respectively.

For the monostable model, there was much less coupling between the IC variations (Figure S1). CaMKII dynamics for peak value and time to peak value only showed dependence on CaMKII IC, and PP1 dynamics were likewise dependent on PP1 IC. Both CaMKII and PP1 peaked very quickly in the monostable model with all peaks by about 300 ms (Figure S2c). As seen in Figure S1c-d, the monostable model produces a single steady state for both CaMKII and PP1 of zero; this is in sharp contrast to the range of steady states seen in the bistable model. While bound AMPAR peak value and steady state appears entirely dependent on CaMKII IC, the time to peak value again showed coupled CaMKII/PP1 IC dependence (Figure S2c), however with the opposite trend as the bistable model. For the monostable model, higher PP1 IC lead to a slower time to peak, while for the bistable model, higher PP1 lead to faster peak time. We note that the small differences within the longer peak value times of the bound AMPAR in the monostable model are small numerical differences.

We also considered these IC variations for knockout trafficking conditions, Figure S3. The no endo/exocytosis case showed similar bound AMPAR dynamics to the control case, while the no influx case showed bistable bound AMPAR behavior for the bistable model and a monostable bound AMPAR for the monostable model, fittingly.

**Figure S2:**
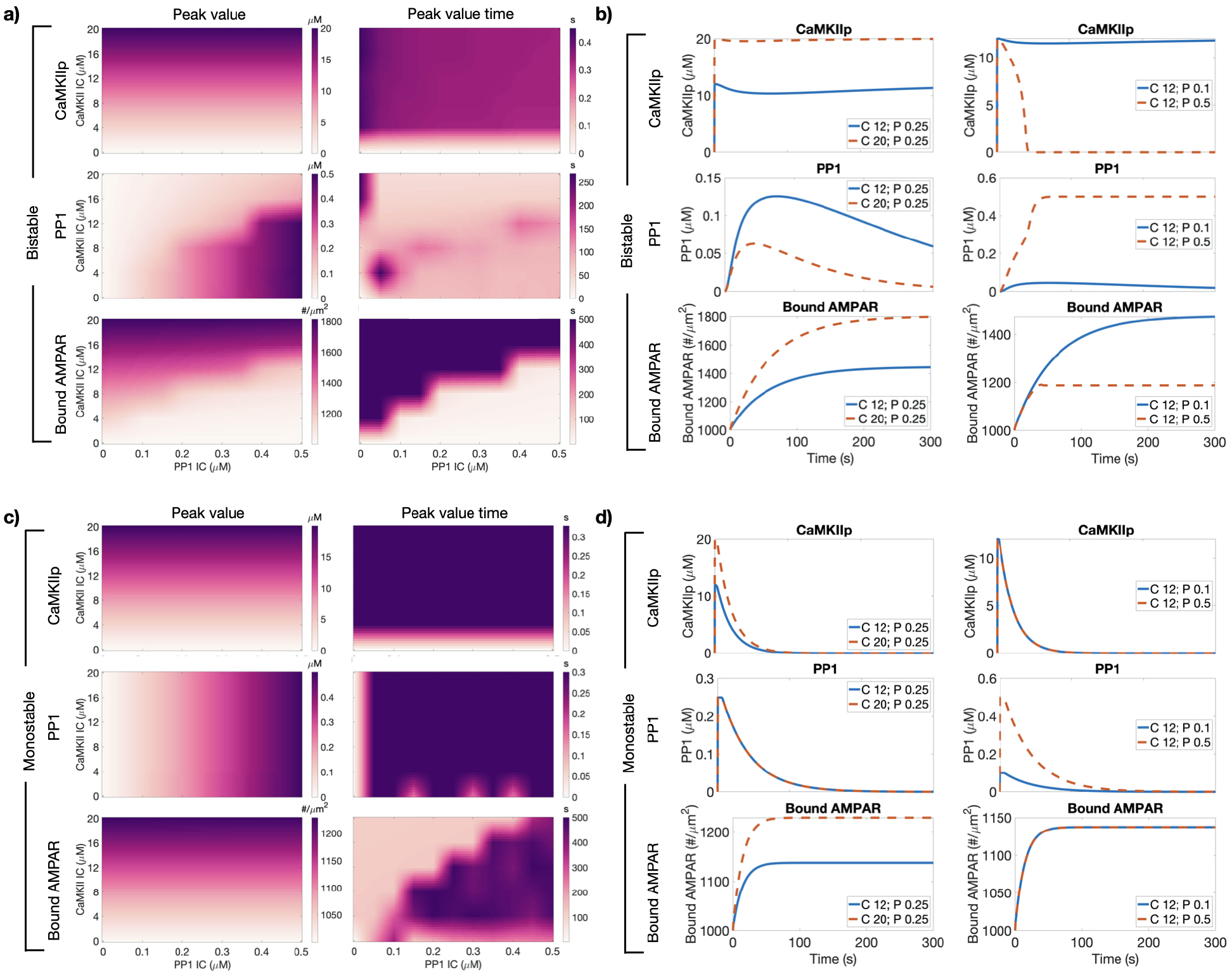
Variations in initial conditions for inactive CaMKII and PP1 influences active CaMKII, active PP1, and bound AMPAR dynamics. a) Peak value and time of peak value (left and right columns respectively) for active CaMKII, active PP1, and bound AMPAR (top to bottom row, respectively) for the bistable model. b) Temporal dynamics for active CaMKII, active PP1, and bound AMPAR (left to right columns, respectively) for a set PP1 IC and varied CaMKII IC (top row) and for a set CaMKII IC and varied PP1 IC (bottom row) for the bistable model. c) Peak value and time of peak value (left and right columns respectively) for active CaMKII, active PP1, and bound AMPAR (top to bottom row, respectively) for the monostable model. d) Temporal dynamics for active CaMKII, active PP1, and bound AMPAR (left to right columns, respectively) for a set PP1 IC and varied CaMKII IC (top row) and for a set CaMKII IC and varied PP1 IC (bottom row) for the monostable model.

**Figure S3:**
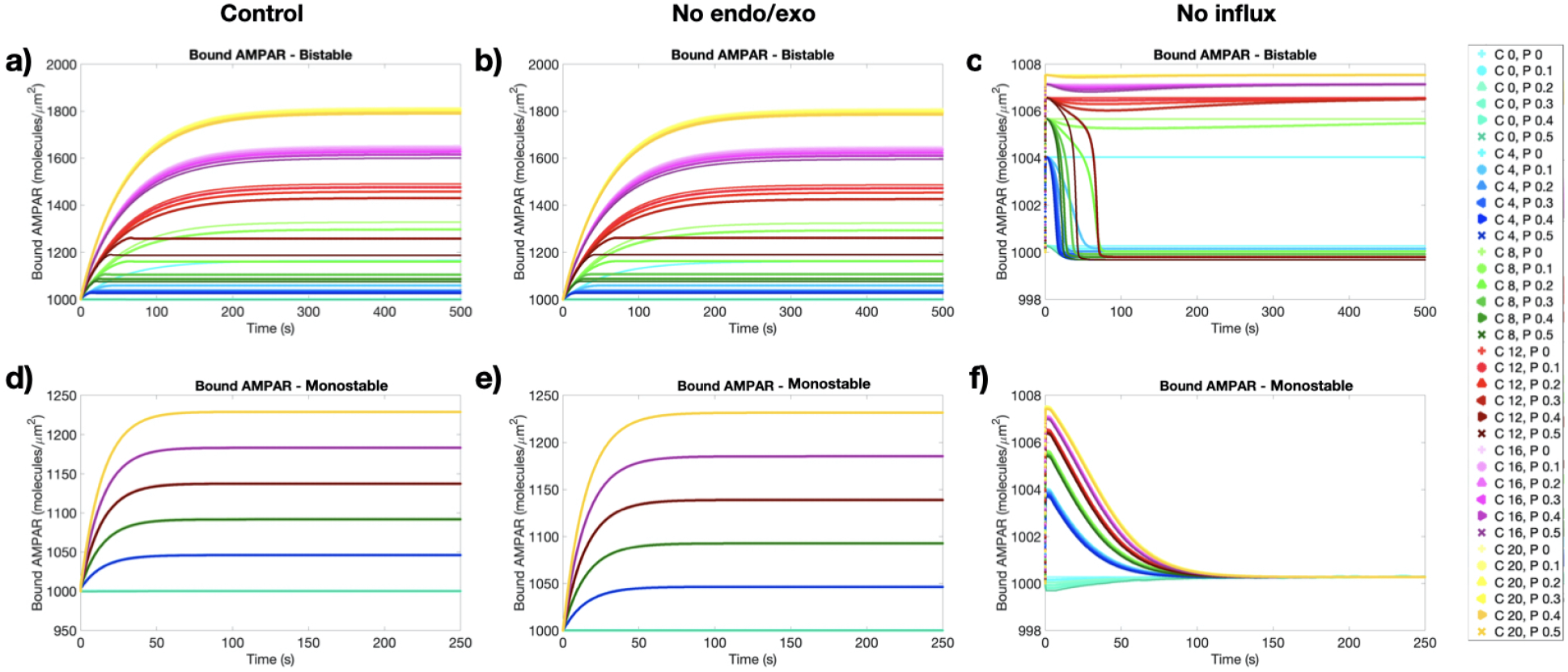
Temporal dynamics for bound AMPAR for both the bistable and monostable models for all CaMKII and PP1 IC combinations for the various trafficking cases. The control case (a), no endo/exocytosis case (b), and no influx case (c) for the bistable model show coupled dependence on the CaMKII/PP1 IC. The control case (d), no endo/exocytosis case (e), and no influx case (f) for the monostable model only show dependence on the CaMKII IC. The C and P in legend stand for CaMKII and PP1, respectively.

**Figure S4:**
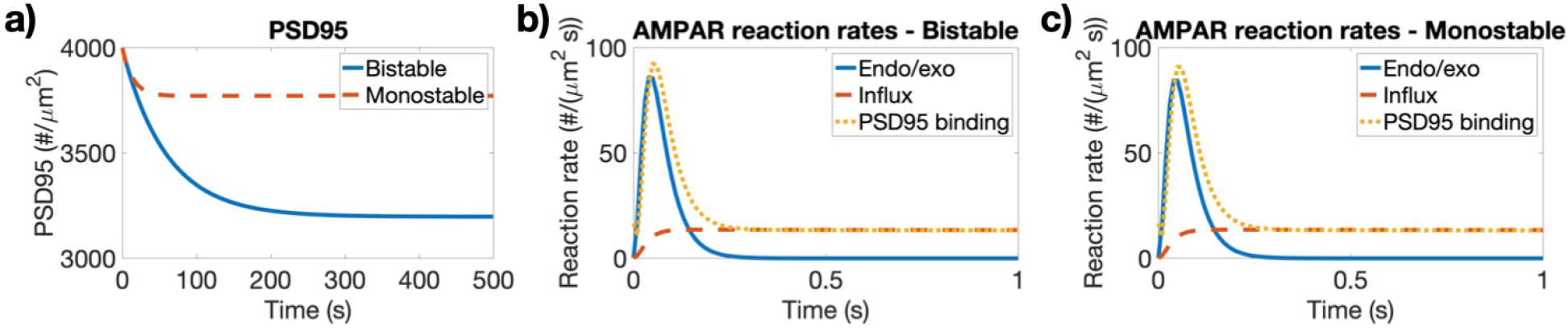
a) Temporal dynamics for PSD95 in the control trafficking case with control IC of 20 μM for CaMKII and 0.25 μM for PP1. Reaction rates for the different trafficking contributions that influence AMPAR dynamics for the bistable (b) and monostable (c) models show similar trends with large rapid PSD95 binding, a smaller endo/exocytosis activity over similar timescales, and then a sustained smaller contribution due to CaMKII-mediated influx.

**Figure S5:**
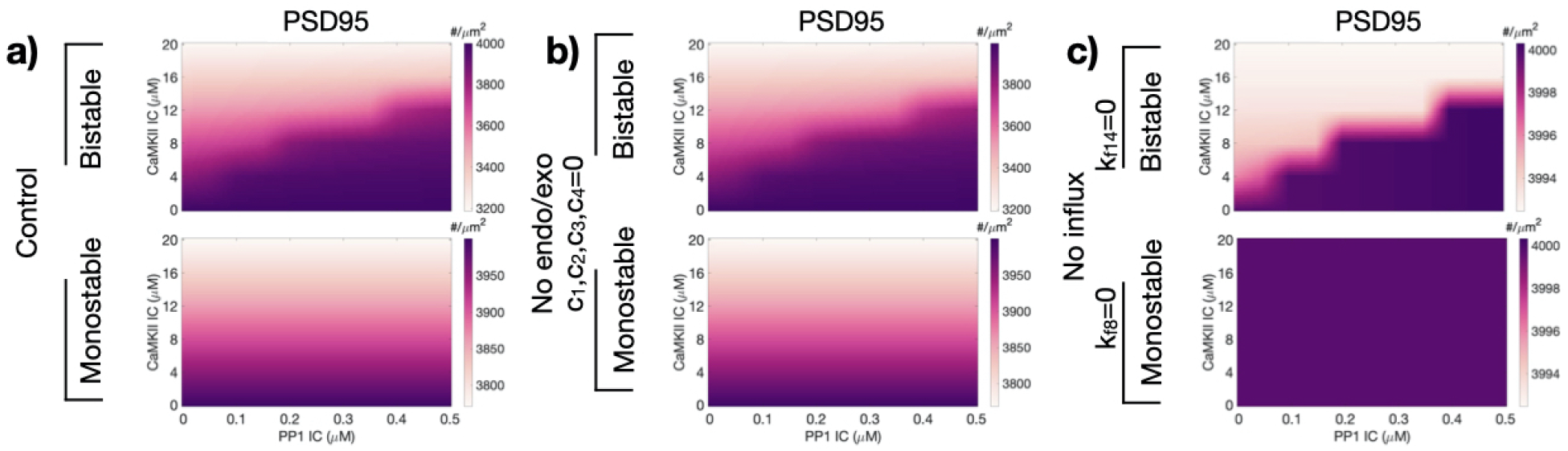
Heatmaps of PSD95 at 500s for a variety of CaMKII and PP1 initial conditions. a) Steady state values for the control trafficking case with all trafficking contributions show dependence on both CaMKII and PP1 IC for the bistable model (top row), but on only CaMKII IC for the monostable model (bottom row). b) Steady state values for the no endo/exocytosis show similar trends as the control case for both the bistable model (top row) and monostable model (bottom row), showing how the CaMKII/PP1 coupling can influence bound AMPAR dynamics despite not directly impacting Amem through endocytosis and exocytosis. c) Steady state values for the no influx case show clear dependence on both CaMKII and PP1 IC for the bistable model (top row), but homogeneous results for the monostable model (bottom row). Without an influx that leads to an increase in membrane AMPAR, the monostable case has a transient decrease in PSD95 but then returns to its initial condition.

### Variations in endo/exocytosis rates

We considered four different cases for both the bistable and monostable models. We considered the model without and with the influx term (with and without mass balance for membrane AMPAR), and with two different initial conditions for CaMKII, 10 and 20 μM, but only one IC for PP1 (0.25 μM). We also considered two methods to vary endocytosis and exocytosis; 1. vary the c_1_ and c_2_ activity dependent terms; and 2. vary the whole endocytosis and exocytosis terms. We considered bound AMPAR for each of these various conditions, see Figure S6, Figure S7, and Figure S8.

**Figure S6:**
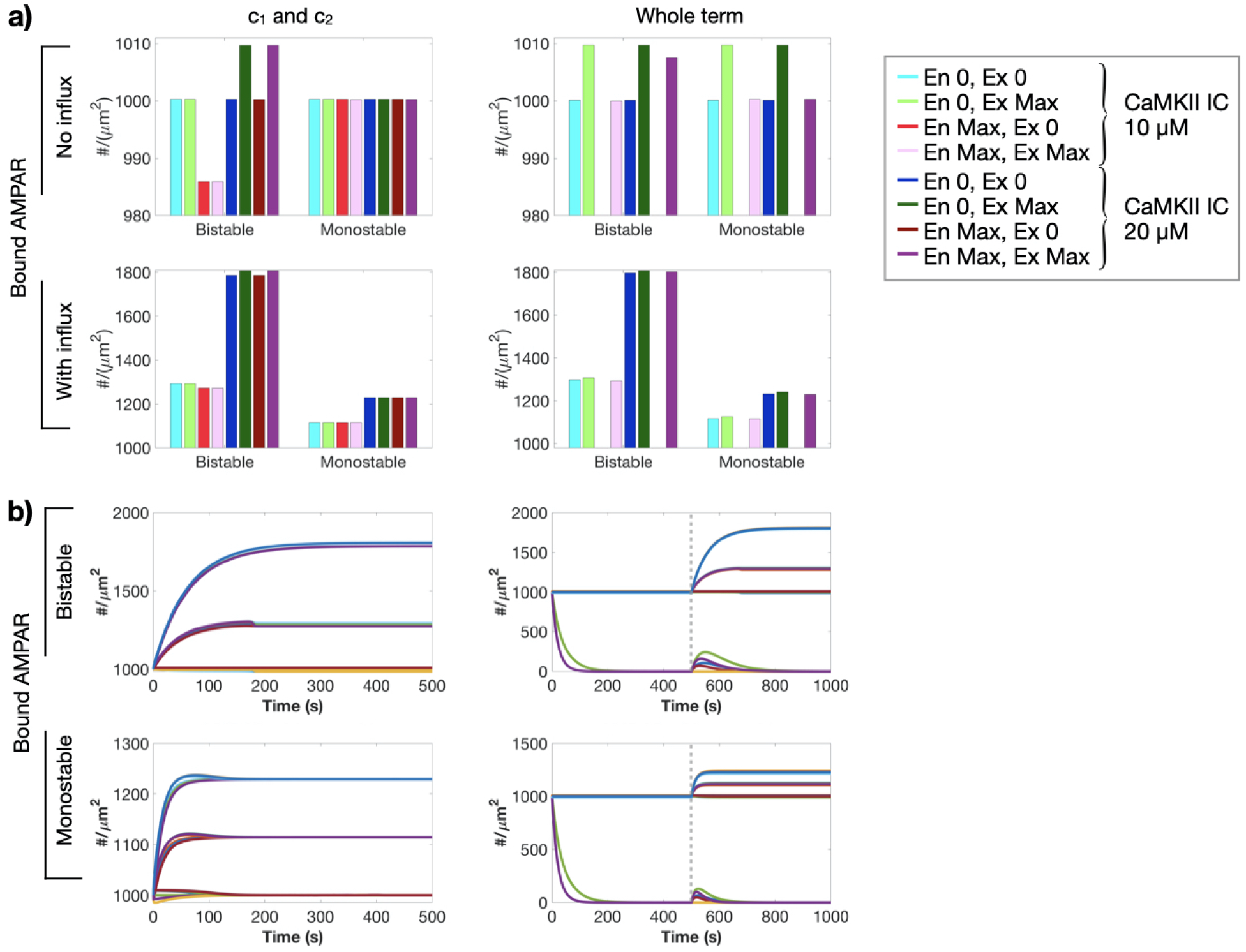
Effect of endocytosis and exocytosis, CaMKII initial condition, and AMPAR influx on bound AMPAR for two different scaling methods. Endocytosis and exocytosis were varied in two different ways: Left column (c_1_ and c_2_): the species dependent rates of c_1_ and c_2_ were scaled; and Right column (Whole term): the whole exocytosis and endocytosis term was scaled. a) Bound AMPAR values at 500 s for the no influx (top row) and with influx (bottom row) cases for different scaling terms of the exocytosis and endocytosis rates (c_1_ and c_2_, respectively) for the two CaMKII ICs for both the bistable and monostable model. We consider two different types of exocytosis and endocytosis scaling, scaling just the c_1_ and c_2_ terms versus scaling the whole endocytosis and exocytosis terms. For the bars that appear vacant, those values are equal to zero (light and dark red bars in the whole term graphs). Gray inset: Legend for panel a where general color corresponds to a set endocytosis (En)/exocytosis (Ex) scaling in this order (En 0, Ex 0; En 0, Ex Max; En Max, Ex 0; En Max, Ex Max), light and dark colors indicate the 10 μM and 20 μM CaMKII IC, respectively. b) Temporal dynamics of bound AMPAR for the bistable (top row) and monostable (bottom row) models for two different CaMKII initial conditions (10 μM and 20 μM), different contributions of endocytosis and exocytosis, and with and without influx.

**Figure S7:**
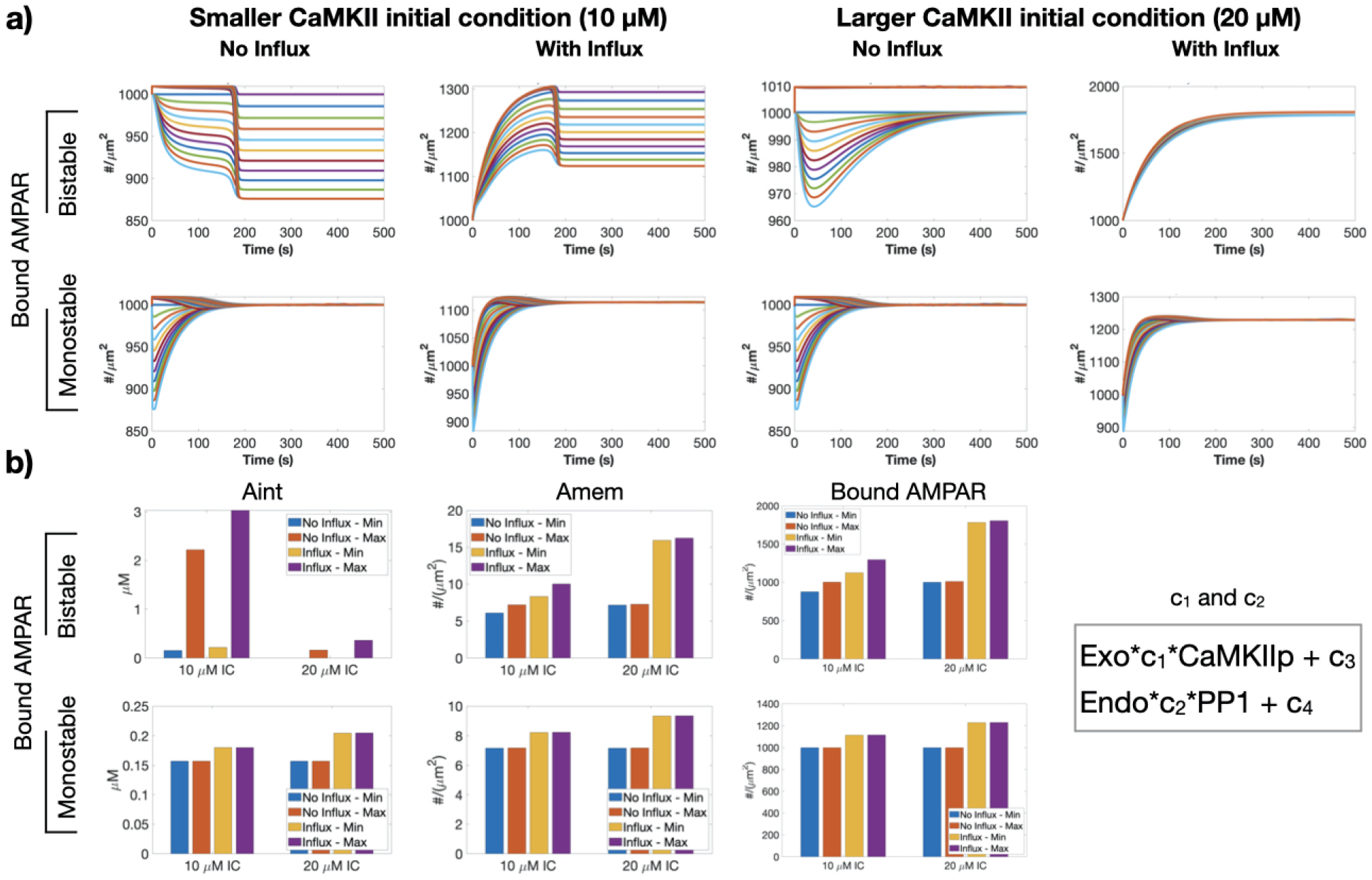
Effect of endocytosis and exocytosis, CaMKII initial condition, and AMPAR influx on bound AMPAR when scaling c_1_ and c_2_. a) Temporal dynamics of bound AMPAR for the bistable (top row) and monostable model (bottom row) for two different CaMKII initial conditions (10 μM, left two columns; 20 μM, right two columns) with and without the membrane AMPAR influx. b) Minimum and maximum bound AMPAR steady states for each CaMKII IC, with and without the influx term for both the bistable and monostable models (top and bottom row, respectively). Gray inset: mathematical representation of how the exocytosis and endocytosis terms are scaled by Exo and Endo.

**Figure S8:**
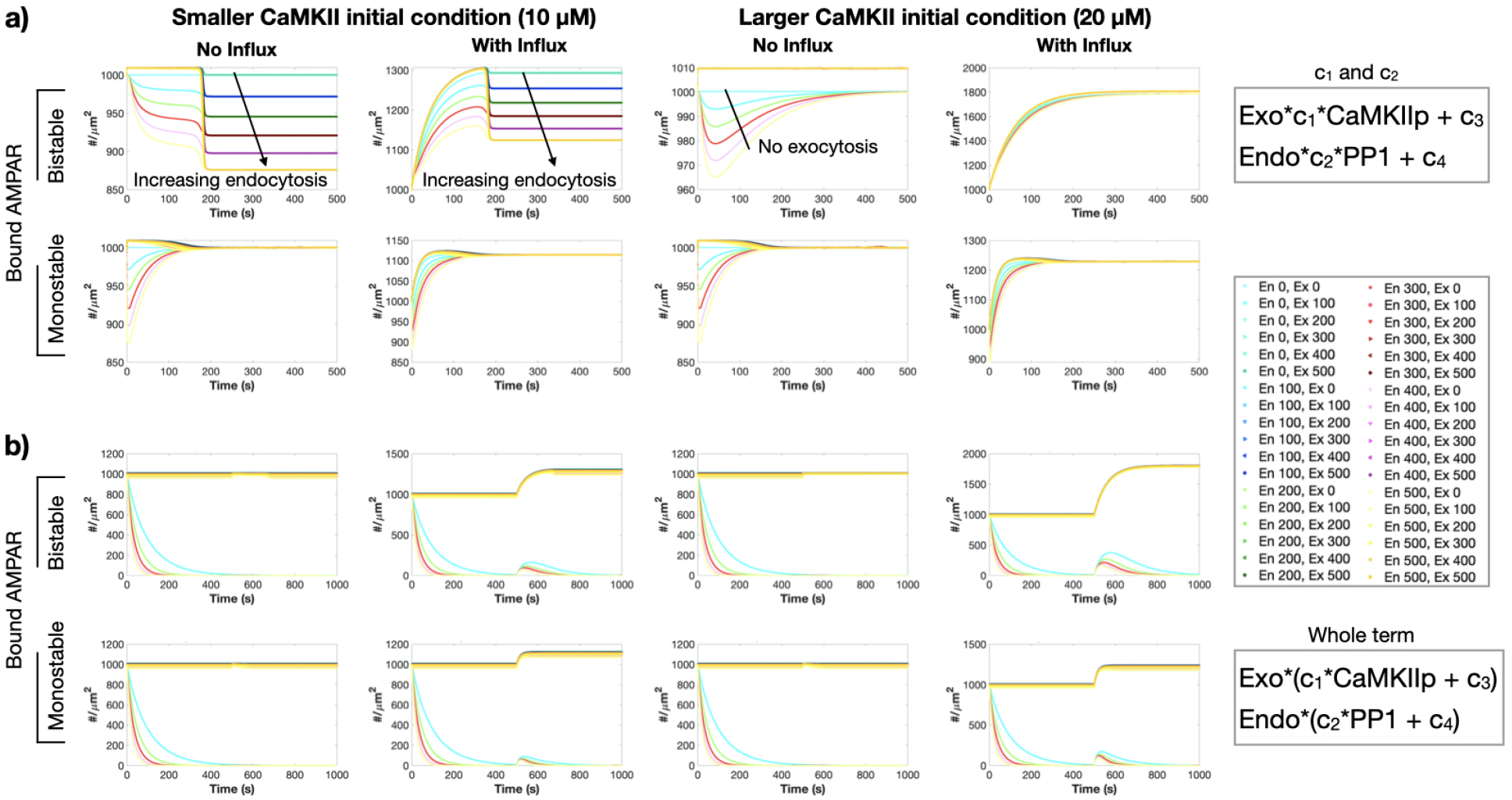
Effect of endocytosis and exocytosis, CaMKII initial condition, and AMPAR influx on bound AMPAR temporal dynamics for two different scaling methods. Endocytosis and exocytosis were varied in two different ways: a) (c_1_ and c_2_): the species dependent rates of c_1_ and c_2_ were scaled (gray inset: mathematical representation); and b) (Whole term): the whole exocytosis and endocytosis terms ere scaled (gray inset: mathematical representation). a) Temporal dynamics of bound AMPAR for the bistable (top row) and monostable model (bottom row) for two different CaMKII initial conditions (10 μM, left two columns; 20 μM, right two columns) with and without the membrane AMPAR influx when varying c_1_ and c_2_. b) Temporal dynamics of bound AMPAR for the bistable (top row) and monostable model (bottom row) for two different CaMKII initial conditions (10 μM, left two columns; 20 μM, right two columns) with and without the membrane AMPAR influx when varying the whole endocytosis and exocytosis terms. Middle gray inset: Legend for panel a. Panel b has the same scaling pattern but the scaling is from 0 to 1 in steps of 0.1. A general color represents a endocytosis scaling, while gradients within that color correspond to exocytosis scalings.

**Figure S9:**
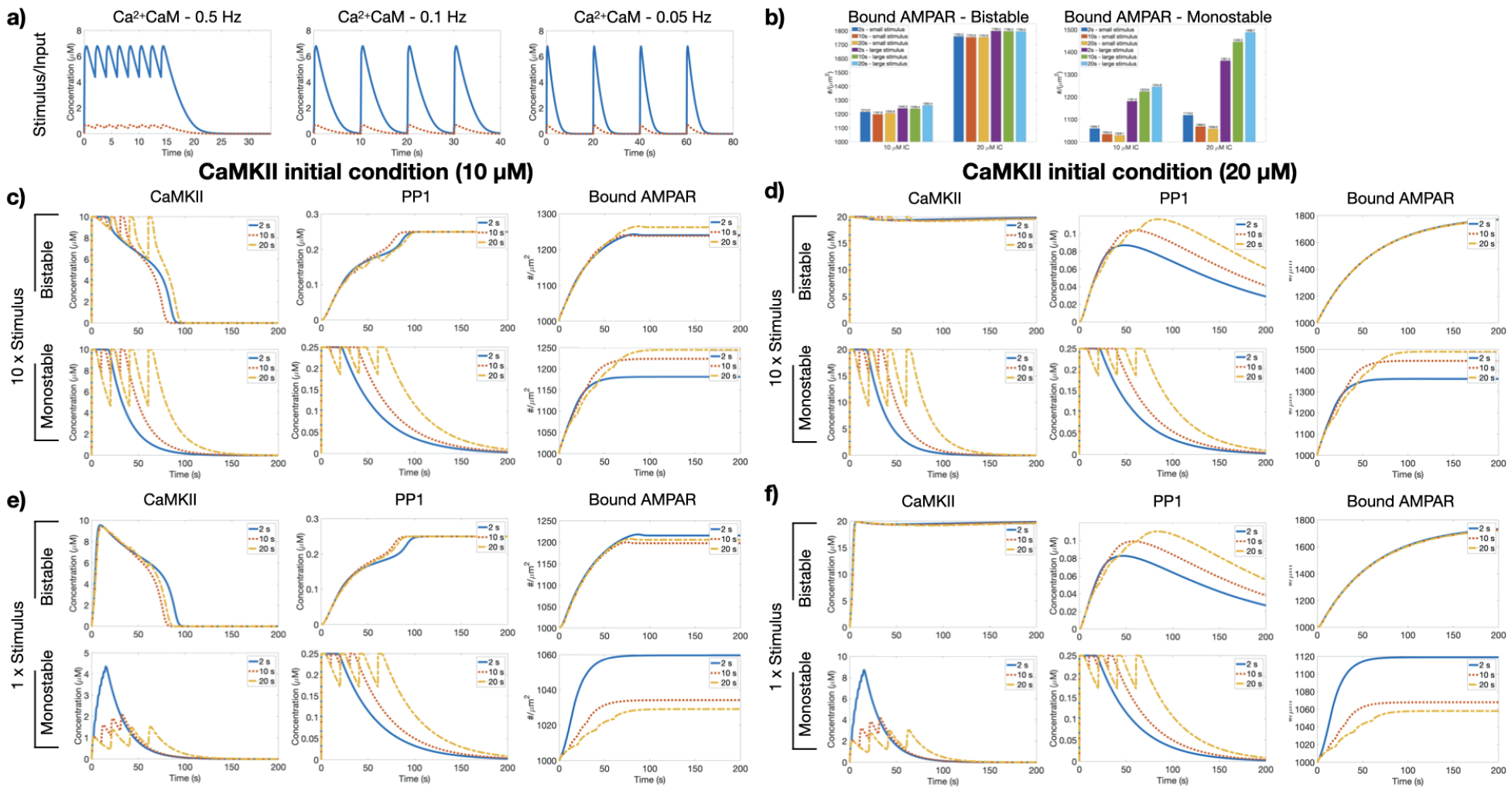
Effect of multiple active Calmodulin spikes on CaMKII, PP1, and bound AMPAR dynamics for different stimulus strengths and CaMKII IC. a) Multiple spikes of active CaM were input into the different model systems at different frequencies (0.5 Hz, 0.1 Hz, 0.05 Hz; left, middle, and right, respectively) at two different amplitudes (large - blue, small - red). b) Steady state values of bound AMPAR for all conditions shown here for both the bistable and monostable models. c) Active CaMKII, active PP1, and bound AMPAR dynamics for both the bistable and monostable model for the large Ca^2+^CaM stimulus and CaMKII IC of 10 μM. d) Active CaMKII, active PP1, and bound AMPAR dynamics for both the bistable and monostable model for the large Ca^2+^CaM stimulus and CaMKII IC of 20 μM. e) Active CaMKII, active PP1, and bound AMPAR dynamics for both the bistable and monostable model for the small Ca^2+^CaM stimulus and CaMKII IC of 10 μM. f) Active CaMKII, active PP1, and bound AMPAR dynamics for both the bistable and monostable model for the small Ca^2+^CaM stimulus and CaMKII IC of 20 μM.

